# Control of Biofilm Formation by an *Agrobacterium tumefaciens* Pterin-Binding Periplasmic Protein Conserved Among Pathogenic Bacteria

**DOI:** 10.1101/2023.11.18.567607

**Authors:** Jennifer L. Greenwich, Justin L. Eagan, Nathan Feirer, Kaleb Boswinkle, George Minasov, Ludmilla Shuvalova, Nicole L. Inniss, Jakka Raghavaiah, Arun K. Ghosh, Karla J.F. Satchell, Kylie D. Allen, Clay Fuqua

**Author notes:** Clay Fuqua, **E-mail:**, Current addresses: J.L.E., Medical Microbiology and Immunology, University of Wisconsin, Madison, WI 53706; N.F., Promega Corp., Madison, WI, 53711. **Author contributions:** J.L.G, J.L.E., K.D.A. and C.F. designed the study; J.L.G., J.L.E., N.F., and K.B. performed the genetic and microbiological experiments, and J.L.G., K.B. and K.D.A. performed biochemical analysis of pterins and pterin-binding. L.S. and G.M. performed crystallization and structural determination. J.R. and A.K.G. developed and implemented a new pterin synthesis method. J.L.G., K.D.A., N.L.I., G.M. K.J.S.F. and C.F wrote the manuscript with input from all authors. C.F. supervised the entire study. **Competing Interest Statement:** K.J.F.S. has a significant interest in Situ Biosciences, a contract research organization that conducts research unrelated to this study. All other authors declare no conflicts of interest. The funders had no role in study design, data collection, and analysis, decision to publish, or preparation of the manuscript.

## Abstract

Biofilm formation and surface attachment in multiple Alphaproteobacteria is driven by unipolar polysaccharide (UPP) adhesins. The pathogen *Agrobacterium tumefaciens* produces a UPP adhesin, which is regulated by the intracellular second messenger cyclic diguanylate monophosphate (cdGMP). Prior studies revealed that DcpA, a diguanylate cyclase-phosphodiesterase (DGC-PDE), is crucial in control of UPP production and surface attachment. DcpA is regulated by PruR, a protein with distant similarity to enzymatic domains known to coordinate the molybdopterin cofactor (MoCo). Pterins are bicyclic nitrogen-rich compounds, several of which are formed via a non-essential branch of the folate biosynthesis pathway, distinct from MoCo. The pterin-binding protein PruR controls DcpA activity, fostering cdGMP breakdown and dampening its synthesis. Pterins are excreted and we report here that PruR associates with these metabolites in the periplasm, promoting interaction with the DcpA periplasmic domain. The pteridine reductase PruA, which reduces specific dihydro-pterin molecules to their tetrahydro forms, imparts control over DcpA activity through PruR. Tetrahydromonapterin preferentially associates with PruR relative to other related pterins, and the PruR-DcpA interaction is decreased in a *pruA* mutant. PruR and DcpA are encoded in an operon that is conserved amongst multiple Proteobacteria including mammalian pathogens. Crystal structures reveal that PruR and several orthologs adopt a conserved fold, with a pterin-specific binding cleft that coordinates the bicyclic pterin ring. These findings define a new pterin-responsive regulatory mechanism that controls biofilm formation and related cdGMP-dependent phenotypes in *A. tumefaciens* and is found in multiple additional bacterial pathogens.

**SIGNIFICANCE:** Biofilms are bacterial communities attached to surfaces, physiologically distinct from free-living cells, and a common cause of persistent infections. Here we define the mechanism of a novel biofilm regulatory system based on excreted metabolites called pterins, that is conserved within a wide range of Gram-negative bacteria, including multiple pathogens of animals and plants. The molecular mechanism of pterin-dependent regulation is reported including structural determination of several members of a new family of pterin-binding proteins. Pterins are produced across all domains of life and mechanistic insights into this regulatory circuit could lead to new advances in antibiofilm treatments.

## INTRODUCTION

The regulation of bacterial attachment to surfaces plays a critical role in the formation of biofilms and can dictate their maturation. Biofilms are surface-associated microbial assemblages that are common among bacteria, and result in dramatic physiological changes including substantial tolerance towards antibiotic treatment (1, 2). Biofilms represent a major challenge for the treatment of bacterial infections as these multicellular structures are recalcitrant to standard therapies and act as protective reservoirs for pathogenic bacteria. Production of bacterial surface structures known as adhesins drive stable attachment of bacteria to surfaces, the first step in biofilm formation (3). Regulation of adhesin elaboration and activity thus can influence where and how biofilm formation occurs. In many bacteria the transition from a free-living to a sessile mode of growth is under the regulatory control of the cytoplasmic second messenger cyclic diguanylate monophosphate (cdGMP). Increasing levels of cdGMP often promote attachment and biofilm formation through production of adhesive proteins and polysaccharides that drive the attachment process (4). Synthesis of cdGMP is catalyzed by diguanylate cyclases (DGCs) and its turnover is driven by phosphodiesterases (PDEs). Single bacterial taxa can have multiple DGCs and PDEs that influence the cdGMP pool, and multi-domain proteins with dual DGC and PDE activities are not uncommon. Environmentally responsive modulation of cdGMP pools is mediated through control of gene expression and through allosteric regulation of these enzymes, many of which have sensory input modules. The flux of this second messenger in cells reflects the combined output of these synthesis and degradation activities (5).

The facultative plant pathogen, *Agrobacterium tumefaciens*, utilizes secreted polysaccharides to stably attach to biotic and abiotic surfaces, most prominently cellulose and a polar adhesin known as the <u>u</u>ni<u>p</u>olar <u>p</u>olysaccharide (UPP) (6). *A. tumefaciens* is the causative agent of crown gall, a plant neoplastic disease that results from bacteria-to-plant horizontal gene transfer (7). Attaching to plant tissue is a requisite step in the virulence pathway of *A. tumefaciens*, but pathogenesis also involves additional plant-produced-signals to activate virulence (8). Production of the UPP, as well as cellulose, is under complex environmental control through cdGMP (9, 10). *A. tumefaciens* encodes close to 30 proteins with predicted DGC domains and only two solo PDE enzymes, but multiple proteins with predicted DGC domains also have PDE domains (6).

In our prior studies, we identified the DcpA protein, with both canonical DGC and PDE domains (9). DcpA has DGC and PDE activity *in vivo*, and genetic evidence suggests that both domains are functional. However, under standard laboratory conditions, DcpA is predominantly a PDE, maintaining low cdGMP levels and thereby limiting surface attachment. The N-terminal portion of DcpA has two transmembrane domains that flank an ∼140 aa periplasmic domain of unknown function. Our prior findings identified several other regulatory components critical for maintaining DcpA as a PDE. These additional regulators include the PruA pteridine reductase, which reduces specific 7,8-dihydropterins to their 5,6,7,8-tetrahydro forms, and PruR, a putative pterin-binding protein (9, 11). Both of these proteins are required to regulate the enzymatic activity of DcpA, and null mutants for *pruA* and *pruR* similarly lead to elevated cdGMP and aberrant activation of surface adhesion. The *pruR* gene is transcriptionally coupled with *dcpA* in a two gene operon, consistent with their related functions.

Pterins are characterized by a nitrogen-rich, bicyclic ring structure with side chains of varying length and modifications extending from the ring carbon at the sixth position (12). Well known biomolecules with a pterin ring include the folates in which the side chain is composed of a para-amino benzoic acid group conjugated to a glutamic acid residue (or poly-glutamate). Folate derivatives are required as cofactors for multiple aspects of one-carbon metabolism, and folate is essential in most organisms (13). More broadly, diverse pterin derivatives are found in all domains of life, and in bacteria are known to act as enzymatic cofactors. Molybdopterin cofactor (MoCo) is a complex molybdenum-containing derivative used by various organisms as a prosthetic group to catalyze redox reactions. MoCo is synthesized from GTP via a pathway independent from folate biosynthesis (14). Biopterin and monapterin are pterins with hydroxylated 3 carbon side chains that function with cytoplasmic amino acid hydroxylases, but curiously the dominant fraction of pterins is detected outside of cells (15). Pterins can exist in a fully oxidized state as well as the reduced 7,8-dihydro and 5,6,7,8-tetrahydro states, the latter of which serves as the biologically active form of the respective cofactor. In *A. tumefaciens,* PruA catalyzes the NADPH-dependent reduction of H_2_MPt to H_4_MPt (11). Extracts of *A. tumefaciens* contain a methylated derivative of H_4_MPt (2′- *O*-methylmonapterin) but this is undetectable in *pruA* deletion mutants and in mutants expressing a catalytically inactive PruA protein (9). Mutations that interfere with PruA activity result in high level DGC activity from DcpA, driving elevated UPP-dependent adhesion.

The regulation of DcpA by PruA is indirect and requires PruR, but the mechanistic basis for this was undefined (9)(Fig. S1A). PruR shares distant amino acid sequence similarity with the SUOX family of proteins, defined by sulfite oxidases and related enzymes, including YedY from *Escherichia coli* (16). SUOX proteins have domains that associate with MoCo to facilitate their enzymatic activities and have a conserved cysteine residue that coordinates the molybdenum atom in MoCo. There is no cysteine at this position in PruR and the protein has no predicted enzymatic activity (9, 17). Rather, we hypothesize that PruR is a pterin-binding protein functioning with non-MoCo pterin derivatives. Here we report on the pterin-response mechanism, the structure of PruR, its control of DcpA, and conservation of this system in multiple bacterial pathogens.

## RESULTS

### PruR regulates DcpA and binds to pterin ligands *in vitro*

The impact of elevated cdGMP on UPP and cellulose production can be qualitatively observed by cultivating *A. tumefaciens* derivatives on solid medium supplemented with the azo-dye Congo Red – increased red pigmentation of colonies (the <u>e</u>levated <u>C</u>ongo <u>R</u>ed or ECR phenotype) is indicative of increased polysaccharide production (10). The ECR phenotype is also predictive for increased surface adhesion via the UPP. The *pruR* gene (ATU_RS16195) is encoded in an operon 9 bp immediately upstream of *dcpA* (ATU_RS16200). Our previous studies revealed that a precise in-frame deletion of the entire *pruR* coding sequence leads to a *dcpA*-dependent ECR phenotype and increased biofilm formation via elevated UPP production (9). We constructed a new deletion mutant, preserving potential translational coupling with *dcpA*, while deleting most of the *pruR* gene. This new Δ*pruR* mutant exhibited a pronounced ECR phenotype and increased biofilm formation, and in contrast to our prior Δ*pruR* mutant, which impacted downstream *dcpA* expression (9), was well complemented with ectopic expression of *pruR* alone (Fig. S1B, S1C).

PruR was purified from *E. coli* to examine pterin binding *in vitro*. A restricted set of pterins are available commercially, and the tetrahydro forms are quite susceptible to oxidation. We synthesized an optically active H_2_MPt from L-xylose by following a related reported procedure (18) and obtained dihydroneopterin (H_2_NPt), dihydrofolate (H_2_F) and tetrahydrofolate (H_4_F) from commercial sources. Our prior studies established that the *A. tumefaciens* pteridine reductase PruA reduces H_2_MPt and H_2_NPt (11). An enzyme-coupled binding assay was developed to actively convert the dihydropterins to their tetrahydro derivatives (Fig. 1A), in which purified pteridine reductase PruA (His_6_-PruA) was incubated with the dihydropterin substrates and NADPH and in the presence of purified non-tagged PruR under anaerobic conditions. Extracted pterins bound to PruR were oxidized and measured by HPLC separation by fluorescence measurement (folate was detected by UV) (Fig. S2). PruR associated with H_2_MPt at detectable levels but bound considerably more of the pterin in the PruA-coupled reaction, indicating increased affinity for the fully reduced H_4_MPt (Fig. 1B *p-*value <0.05 compared to all other pterins). PruR also weakly associated with the neopterin derivatives but did not show a statistically significant preference for H_4_NPt compared to H_2_NPt. PruR bound even less well to H_2_F, with no preference for H_4_F; both folate compounds were bound significantly less well than H_4_MPt and H_4_NPt.

**Figure 1.**
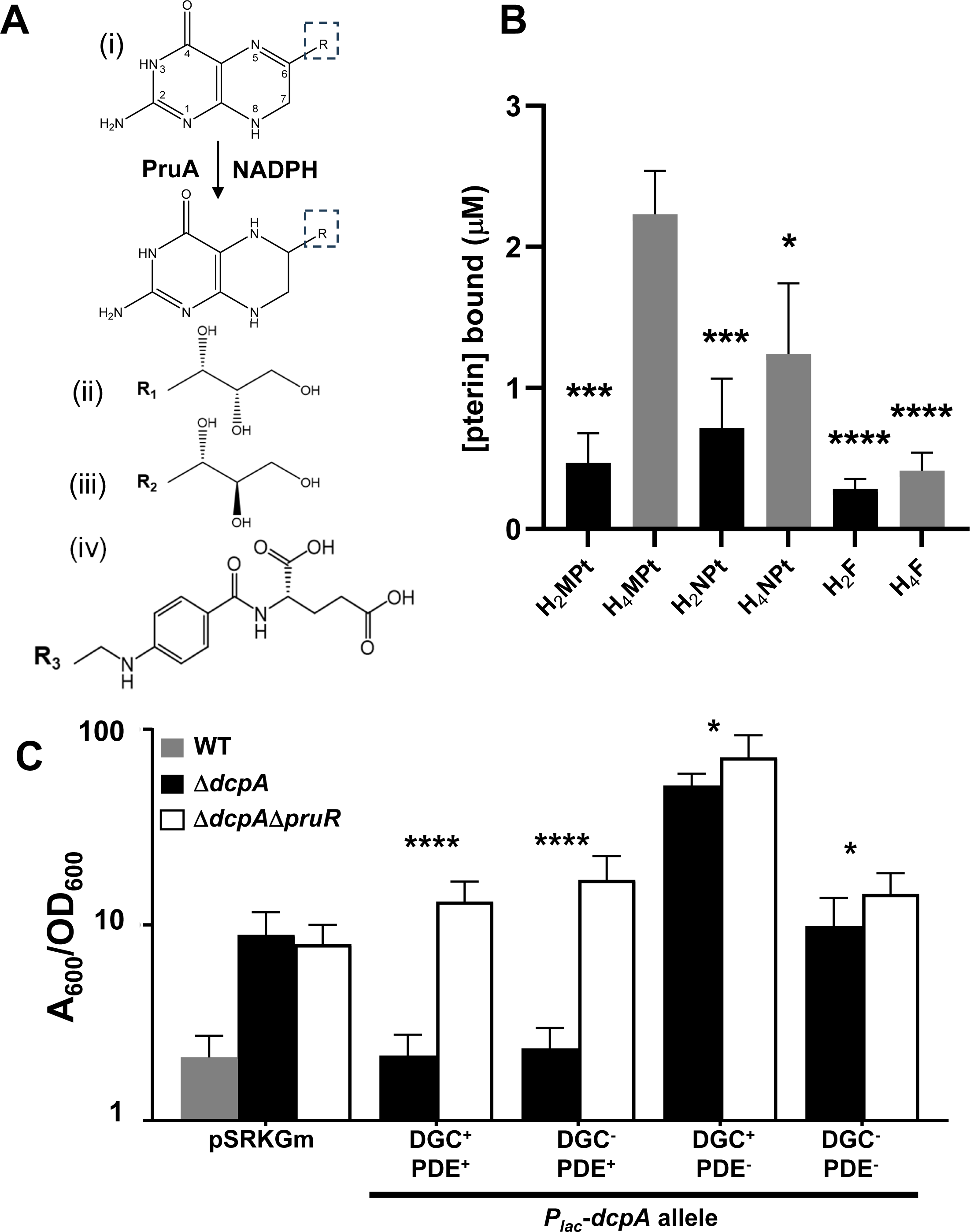
PruR binds pterins and is required to control both the DGC and PDE activity of DcpA. **(A)** PruA reaction and relevant pterin molecule structures. (i): PruA uses NADPH as a cofactor and catalyzes the reduction of a 7,8-dihydropterin substrate to a 5,6,7.8-tetrahydropterin. Atoms are numbered in the dihydropterin. R indicates side groups as shown in (ii)-(iv); (ii): R1-monapterin; (iii): R2-neopterin; (iv): R3 - *p*ABA-glutamate (folate). **(B)** *In vitro* pterin binding assays were performed as described in the supplemental methods with purified Δ_SS_-PruR (50 µM), NADPH, and with or without His_6_-PruA. HPLC fractioned reactions were examined by fluorescence for the oxidized pterins (excitation: 356 nm, emission: 450 nm). UV absorbance at 283 nm was used to measure folate relative to standards. PruR was incubated with the following pterin or folate species: H_2_MPt, dihydromonapterin; H_4_MPt, PruA-generated-tetrahydromonapterin; H_2_NPt, dihydroneopterin; H_4_NPt, PruA-generated tetrahydroneopterin; H_2_F, dihydrofolate; and H_4_F, tetrahydrofolate. Bars are averages of triplicate assays with error bars as standard deviations. Analyzed by standard one-way ANOVA and post-hoc Tukey analysis (*P* values relative to H_4_MPt, *, <0.05, *** <0.001, ****, <0.0001). **(C)** Biofilm assays of. *A. tumefaciens* C58 WT (gray bar), a Δ*dcpA* mutant (black bars) or a Δ*dcpA*Δ*pruR* double mutant (white bars) containing the vector control or a plasmid-borne *P_lac_*-*dcpA* fusion expressing either the wild type *dcpA* or catalytic site mutants (DGC^-^, E308A; PDE^-^, E431A; DGC^-^PDE^-^, both). Ratio of acetic acid-solubilized CV absorbance (A_600_) from 48 h biofilm assays normalized to the OD_600_ planktonic turbidity from the same culture. Assays performed in triplicate and error bars are standard deviation; *P* values calculated comparing complementation of *dcpA* in the Δ*dcpA* strain compared to the Δ*dcpA*Δ*pruR* strain by standard two-tailed *t*-test. *(P* values, * <0.05, **** <0.0001.)

### PruR actively modulates the DGC and PDE activity of DcpA

The *in vivo* phenotypes of several *dcpA* point mutants suggested that it has both DGC and PDE activity, but that the PDE activity is dominant in planktonic laboratory culture (9, 10). Enzymatic assays of the purified DcpA cytoplasmic domain (res. 190-644, see Methods) revealed that the protein has both activities (Fig. S3), although the DGC activity is weaker (0.26 A_360_ units min^-1^mol^-1^) compared to PDE activity (15.7 A_360_ units min^-1^ mol^-1^). However, it was unclear whether the PruR-dependent PDE-dominant activity of DcpA *in vivo* reflects PruR stimulation of the PDE activity, the inhibition of DGC activity, or both.

Ectopic expression of *dcpA* alone in *E. coli*, in the absence of *pruR*, resulted in high level cdGMP synthesis and mutation of the DGC catalytic motif (GGDEF>GGDAF; E308A) abolished this cdGMP increase, suggesting that PruR is required for dominant PDE activity (9). Plasmid-borne expression of wild type *dcpA* complements the elevated biofilm formation of *the* Δ*dcpA* mutant to normal levels. By contrast, the Δ*pruR*Δ*dcpA* mutant harboring this plasmid exhibits a dramatic increase in biofilm formation relative to wild type, and higher than the single *dcpA* mutant (Fig. 1C), consistent with the increased cdGMP caused by *dcpA* ectopic expression in a *pruR* null mutant (9). Expression of the DGC^-^(PDE^+^) DcpA_E308A_ allele in the Δ*dcpA* mutant decreases biofilm formation to wild type levels. However, this allele fails to cause this decrease in the Δ*pruR*Δ*dcpA* mutant, suggesting that PDE activity is under PruR control. Ectopic expression of *dcpA* in wild type does not significantly diminish biofilm formation (likely due to cdGMP production by other DGCs) (9).

In the absence of PruR, the PDE activity of DcpA is not increased by ectopic expression of DcpA. Mutation of *dcpA* to abolish DcpA PDE activity (EAL>EEL; E431A; PDE^-^DGC^+^) resulted in striking stimulation of biofilm formation when this mutant is ectopically expressed in the Δ*dcpA* mutant (Fig. 1C). However, expression of the DcpA_E431A_ allele in the Δ*pruR*Δ*dcpA* mutant imparts even more dramatic biofilm stimulation, suggesting that PruR dampens DGC activity in addition to stimulating PDE activity, and its effects on these two activities are genetically separable. Ectopically expressing an allele mutated for both domains (DGC^-^PDE^-^, E308A E431A,) does not strongly impact biofilm formation in either the Δ*dcpA* or the Δ*pruR*Δ*dcpA* mutant but shows a slight increase in the *pruR-dcpA* double mutant relative to the Δ*dcpA* mutant.

### PruR is secreted and active in the periplasm

Sequence analysis suggests that the PruR protein has an N-terminal secretion signal (aa 1-22, Signal-P score 0.31, Fig. 2A). Given our hypothesis that PruR is a pterin-binding protein, this prediction is surprising, as the only established function of bacterial pterins is to serve as cofactors for amino acid hydroxylases, which are cytoplasmic enzymes (15). To experimentally evaluate whether PruR is secreted to the periplasm *in vivo*, we first tested this genetically by fusing it to *phoA*, encoding the enzyme alkaline phosphatase (AP) that requires localization to the periplasm for activity (19). Plasmid-borne, ectopic expression of the *pruR-phoA* fusion in *A. tumefaciens* led to detectable AP activity in whole cells, above the very low levels of the same strain with no *phoA* (Fig. S4A). Furthermore, fusion of only the *pruR* signal sequence (res. 1-22) with *phoA* (*pruR*_SS_*-phoA*) similarly expressed from the same plasmid resulted in much stronger AP activity. Neither construct was affected by expression in the Δ*pruR* null mutant. It is likely that the full length PruR-PhoA fusion partially diminishes AP activity.

**Figure 2.**
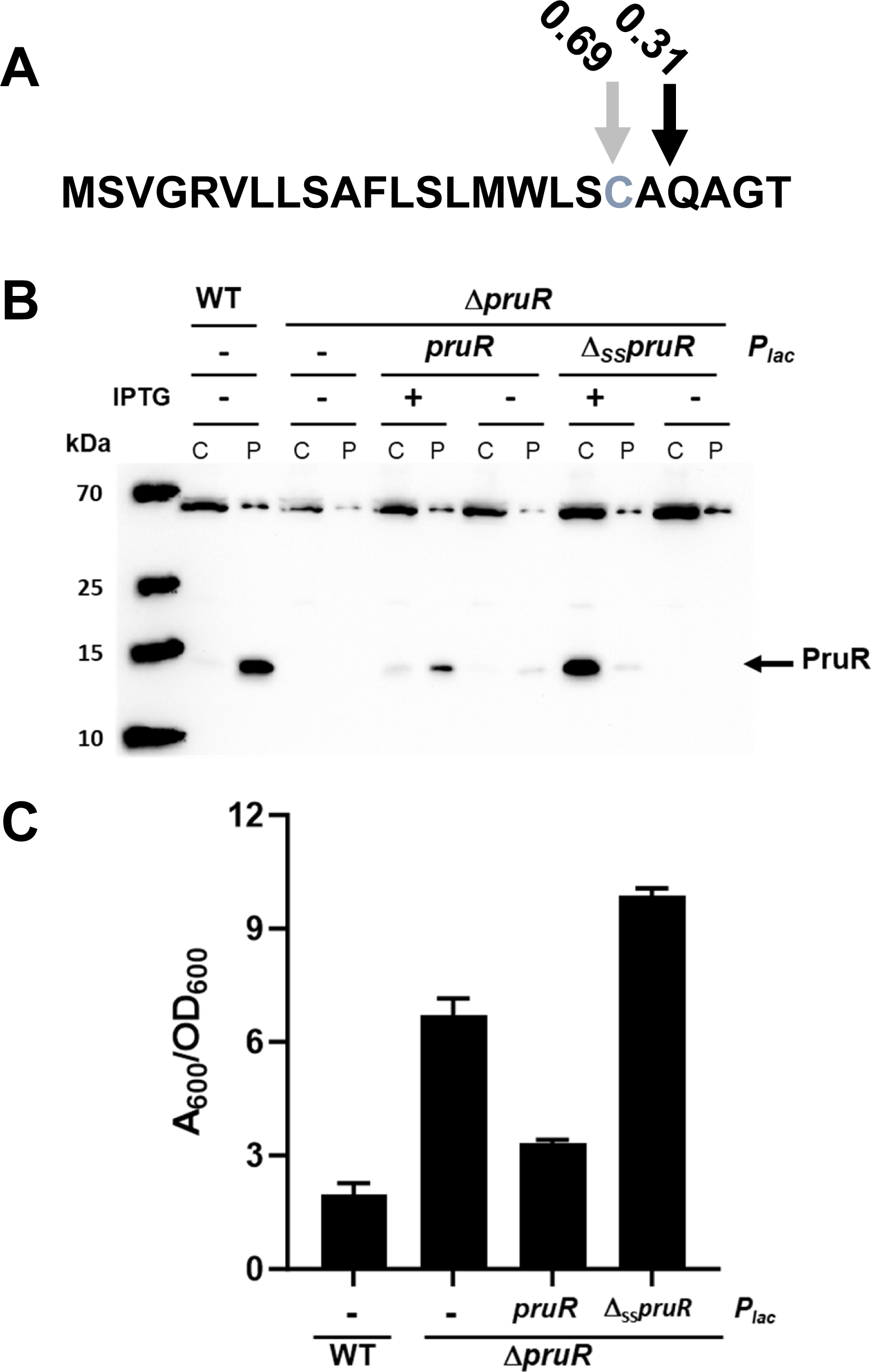
PruR is a periplasmic protein. **(A)** Signal P prediction of the PruR N-terminal signal sequence. Black arrow, predicted signal peptidase cleavage site; Grey arrow, putative cleavage and lipidation site; Grey text, predicted lipidated Cys19. **(B)** Western blot of SDS-PAGE using α-PruR polyclonal antisera to probe extracts of wild type *A. tumefaciens* C58 WT and the Δ*pruR* mutant on its own or expressing either the *P_lac_-pruR* plasmid or the *P_lac_-*Δ*_SS_pruR* plasmid (both plasmids also expressing *dcpA*). Cultures were grown to similar densities with or without induction with 400 μM IPTG and fractionated to separate the cytoplasmic/membrane fraction (C lanes) from the periplasmic fraction (P lanes). **(C)** Biofilm assays of WT or a Δ*pruR* mutant derivative with the empty vector plasmid (-) or harboring a plasmid-borne *P*_lac_ fusion expressing either *pruR* or the Δ_SS_*pruR* (both plasmids also express *dcpA*). Ratio of acetic acid-solubilized CV A_600_ from 48 h biofilm assays normalized to the OD_600_ planktonic turbidity of the same culture. Assays performed in triplicate and error bars are standard deviation; *P* values calculated by standard two-tailed *t*-test (*P* values, *** <0.01).

The periplasmic localization of PruR was tested directly by fractionating *A. tumefaciens* cells from wild type and several derivatives into periplasmic and cytoplasmic fractions using an osmotic shock protocol and probing western blots with a polyclonal antibody preparation raised against PruR (α−PruR). Wild type cells clearly revealed a protein the size of the processed from of PruR (146 aa, predicted 16.1 kDa) in the periplasmic fraction, whereas a Δ*pruR* mutant lacked this protein (Fig. 2B). Ectopic expression of a plasmid-borne copy of *pruR* expressed from *P_lac_* in the Δ*pruR* mutant revealed an IPTG-inducible PruR protein in the periplasmic fraction. In contrast the same expression construct deleted for the *pruR* signal sequence (Δ_SS_*pruR*) results in PruR that remains in the cytoplasmic fraction.

Complementation of a Δ*pruR* mutant (the mutation which is partially polar on *dcpA*) with a plasmid expressing *pruR* and *dcpA* rescues this mutant to normal levels of surface attachment (Fig. 2C). However, expression of the same plasmid expressing the Δ_ss_*pruR* allele and *dcpA* failed to rescue these phenotypes (Fig. 2C). The even greater surface adherence observed with the (*P_lac_*-Δ_ss_*pruR*-*dcpA*) plasmid was most likely due to its additional copy of *dcpA*.

Although PruR has a match for an N-terminal signal sequence there was an even stronger match identified for a lipidation site in the signal sequence predicted at the cysteine 19 (C19, Signal-P score 0.68, Fig. 2A) (20). In this analysis, PruR would be lipidated at C19 with signal peptidase cleavage between it and the adjacent serine (S18, Fig. 2A), rather than A22 as predicted for the non-lipidated secretion signal. Lipidated PruR would be predicted to associate with the periplasmic leaflet of the inner membrane rather than the outer membrane (21). Although this predicted lipidation site is an imperfect match to the spacing for the canonical sequence “lipobox” motif (L[A/S][A/G]C), we tested whether C19 is required for PruR activity. Ectopic expression of a *pruR* allele with this cysteine residue mutated to an alanine (C19A) fully complemented the elevated biofilm phenotype Δ*pruR* mutant, suggesting that PruR is not lipidated, or minimally that it does not have to be anchored to the inner membrane to function in the periplasm (Fig. S4B). Furthermore, *pruR* was effectively secreted and retained full activity when fused with the *malE* and *dsbA* signal sequences (MalE_SS_ and DsbA_SS_) from *E. coli*, well characterized to direct two distinct Sec-dependent secretion mechanisms, and to be non-lipidated (22, 23) (Fig. S4C and S4D).

### *In vivo* crosslinking reveals a PruR-DcpA complex

Direct interaction of PruR with the periplasmic region of DcpA would be one mechanism by which pterins could regulate DcpA DGC and PDE activity. To test this hypothesis, we used disuccinimidyl suberate (DSS) to perform protein crosslinking with cell suspensions of *A. tumefaciens* and then probed for PruR and DcpA proteins using polyclonal antibodies to PruR and separately to the periplasmic portion of DcpA (Fig. 3). PruR is an approximately 16 kDa protein, and full-length DcpA is approximately 70 kDa. Upon DSS addition in wild type cells, SDS-PAGE separation, and western blotting with α-PruR antibody, a new protein species of ∼85 kDa is observed (Fig. 3A). This species is absent in the Δ*dcpA* mutant but is significantly more pronounced in wild type *A. tumefaciens* expressing the *P_lac_*-*pruR*-*dcpA* construct. Elevated expression of the *pruA* gene had no obvious impact on crosslinking of PruR and DcpA. Probing these western blots with a lower titer α-DcpA antibody preparation against the periplasmic domain captured the complex in cells harboring the *P_lac_*-*pruR*-*dcpA* plasmid but was not sufficiently sensitive to detect the complex at native levels of expression (Fig. S5).

**Figure 3.**
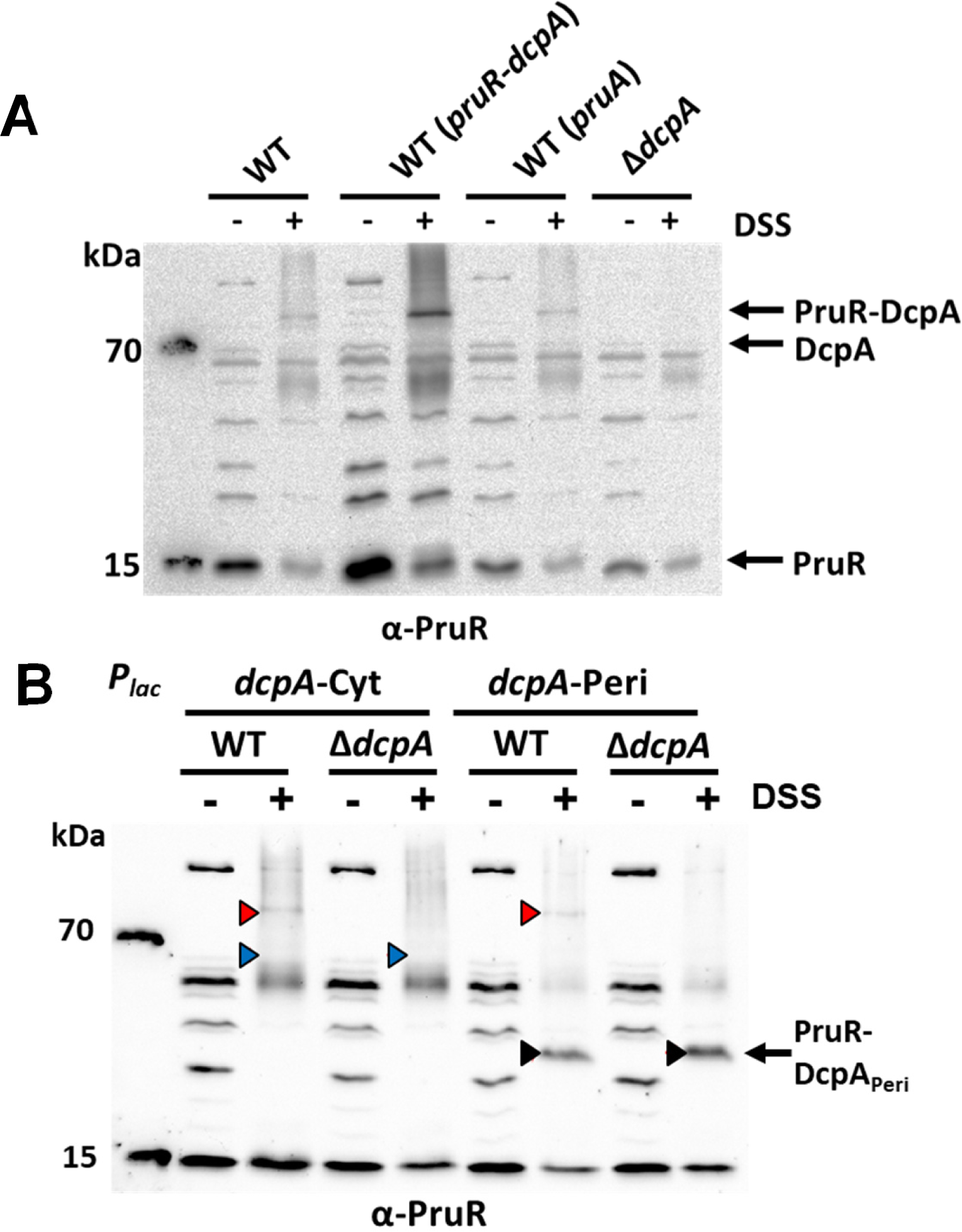
PruR forms a complex with the periplasmic region of DcpA. **(A)** Western blot probing for PruR with α−PruR polyclonal antisera. Whole cell suspensions were either untreated or incubated with DSS crosslinker (0.75 mM) for wild type *A. tumefaciens* C58 and the Δ*dcpA* mutant. Wild type *A. tumefaciens* harboring a plasmid ectopically expressing either *pruR-dcpA* or *pruA* from *P_lac_*(400 μM IPTG) was also subjected to same analysis. Black arrows; PruR-DcpA complex, 87 kDa; DcpA, 71 kDa; PruR; 16 kDa. **(B)** Similar western blot to panel A probing for PruR following DSS crosslinking of whole cell suspensions of either *A. tumefaciens* wild type strain or the Δ*dcpA* mutant. Either the cytoplasmic (DcpA_cyt_) or periplasmic (DcpA_peri_) domains of DcpA are ectopically expressed from *P_lac_* (400 μM IPTG). Red triangles, PruR-DcpA, 87 kDa; Blue triangles, expected size of PruR-DcpA_cyt_, ∼68 kDa; Black triangles, PruR-DcpA_peri_,38 kDa. α-PruR used at 1:40,000 and antibody binding detected with GAR-HRP secondary antibody and chemiluminescent substrate on a BioRad ChemiDoc. Non-specific bands serve as protein loading controls.

Next, we tested the interaction of PruR independently with the periplasmic and cytoplasmic DcpA domains. The DcpA periplasmic domain including its two transmembrane elements (expressing codons 1-192) and the DcpA cytoplasmic expression plasmid (codons 190-644) were independently expressed from *P_lac_* either in the wild type or the Δ*dcpA* mutant. DSS crosslinking of whole cell suspensions revealed efficient PruR crosslinking with the DcpA periplasmic domain (predicted ∼37.4 kDa crosslinked complex) but not with the cytoplasmic domain (predicted 66.5 kDa complex) (Fig. 3B). In the wild type background, the full length PruR-DcpA complex was also visible, when probed for either PruR or the periplasmic domain of DcpA, but this was abolished in the Δ*dcpA* mutant. These results are consistent with the genetic evidence that the PruR-DcpA interaction occurs in the periplasm.

### PruR interaction with the DcpA periplasmic domain is decreased in a *pruA* mutant

The Δ*pruA* mutant manifests a dramatic increase in UPP production and surface attachment, similar to mutants of *pruR* or *dcpA*, and its impact is dependent on the presence of a functional *dcpA* gene (9). To evaluate whether PruA-generated H_4_MPt impacts PruR-DcpA interactions, we compared DSS crosslinking for whole cell suspensions of the Δ*pruA* mutant compared to the wild type, both ectopically expressing PruR and the periplasmic domain of DcpA (these derivatives retain the chromosomal *pruR-dcpA* genes). The amount of complex formation between PruR and DcpA_Peri_ was substantially diminished in the Δ*pruA* mutant relative to the wild type using both the α−PruR (Fig. 4A) and α−DcpA_Peri_ (Fig. 4B) antibodies. These results suggest that *pruA* is required for maximal complex formation between PruR and DcpA and suggests that pterins formed by PruA are essential for the PruR and DcpA interaction.

**Figure 4.**
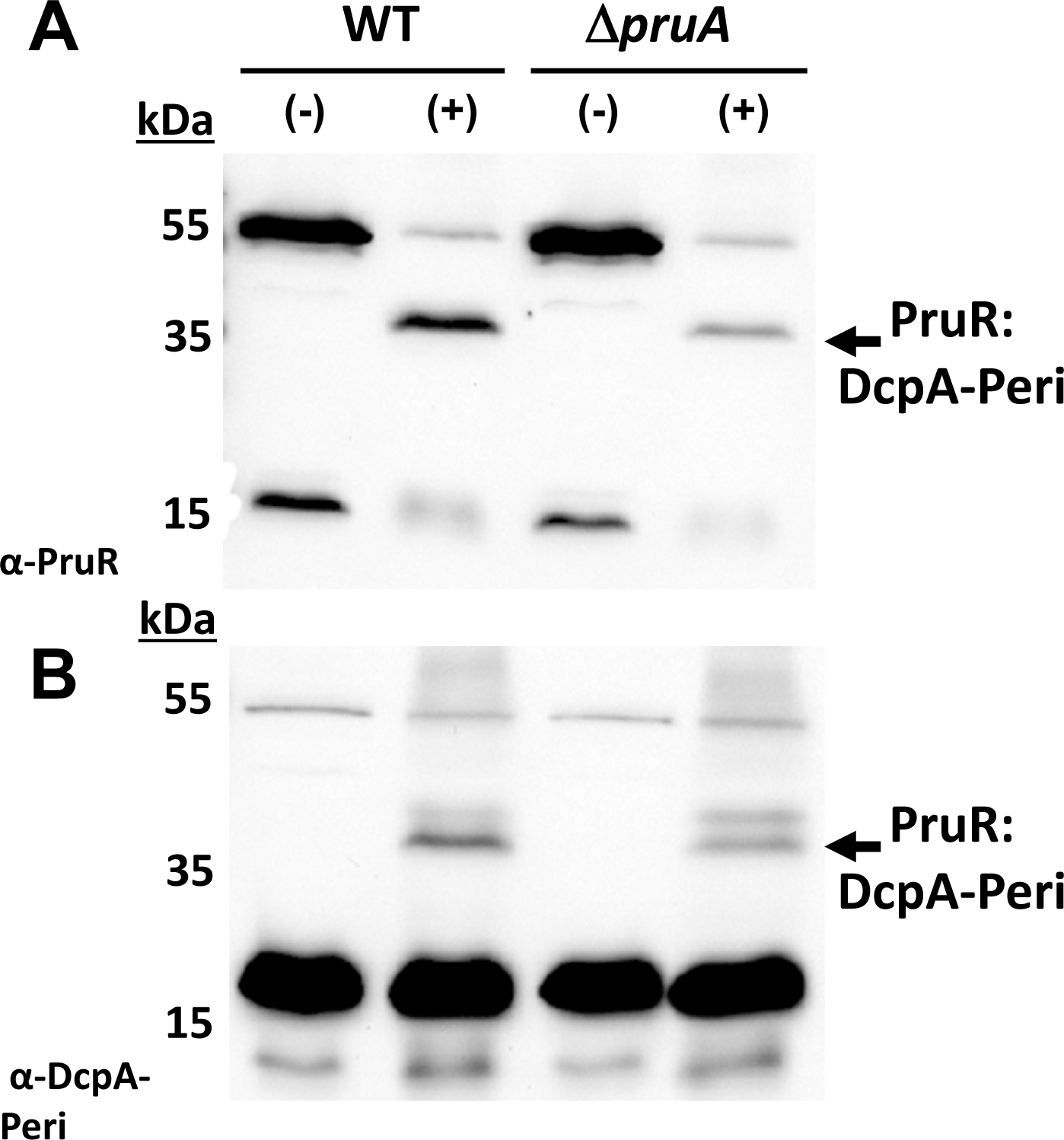
Deletion of *pruA* diminishes the interaction between DcpA and PruR. Western blot probing for the PruR-DcpA_peri_ complex following *in vivo* DSS crosslinking in *A. tumefaciens* wild type or a Δ*pruA* mutant expressing the *P_lac_*-*pruR-dcpA_cyt_* plasmid (400 μM IPTG). Probed with polyclonal antibody preparations **(A)** α-PruR, 1:40,000; **(B)** α-DcpA_peri_, 1:20,000. Antibody binding was detected with GAR-HRP secondary antibody and a chemiluminescent substrate on a BioRad ChemiDoc. Non-specific bands serve as protein loading controls.

### Conservation of *pruR-dcpA* type operons among proteobacterial pathogens

We next sought to determine whether PruR is conserved in other bacteria. There are well-conserved homologs of PruR in several taxa within the Alpha- and Gamma-proteobacteria (Fig. 5). All of these proteins are of similar sizes with an N-terminal secretion signal and share residues common among the SUOX family of proteins (see residues marked by asterisks in Fig. S6A). Notably, all lack the critical cysteine involved in MoCo binding in SUOX proteins, and instead have a tryptophan. None of these has a Cys in their N-terminal signal sequence consistent with our finding that this residue is dispensable for PruR function (Fig. S6A). Additionally, inspection of the genomic location for the PruR homologs in these proteobacteria reveal presumptive bicistronic operons with a downstream gene encoding a DcpA homolog which is comprised of two transmembrane domains flanking a periplasmic loop and a large cytoplasmic portion of the protein. Many of these have both DGC and PDE domains such as DcpA, although in some cases they have only the DGC domain without the C-terminal PDE (Fig. 5). The predicted periplasmic domains of these *dcpA* homologs are roughly the same length (138-150 aa) and are chemically similar. However, these regions are only weakly conserved at the primary amino acid sequence level, with two invariant residues in common, a tryptophan and a glutamate (W40 and E48 in the full-length *A. tumefaciens* DcpA) (Fig. S6B). Strikingly, most of the bacteria that encode this *pruR-dcpA* operon are opportunistic pathogens of animals or plants. Interestingly, *Vibrio cholerae* has two *pruR-dcpA* operons, VC1933-1934 and VCA0075-0074 (Fig. 5). The VC1934 DcpA homolog has both DGC and PDE domains, but the GGDEF motif is degenerate (GADEF) and likely inactive. Conversely, the VCA0074 DcpA homolog only has the GGDEF domain without the PDE domain. In fact, VCA0074 is the well-studied CdgA protein established to have DGC activity in *V. cholerae* and to play an important role in biofilm formation (24).

**Figure 5.**
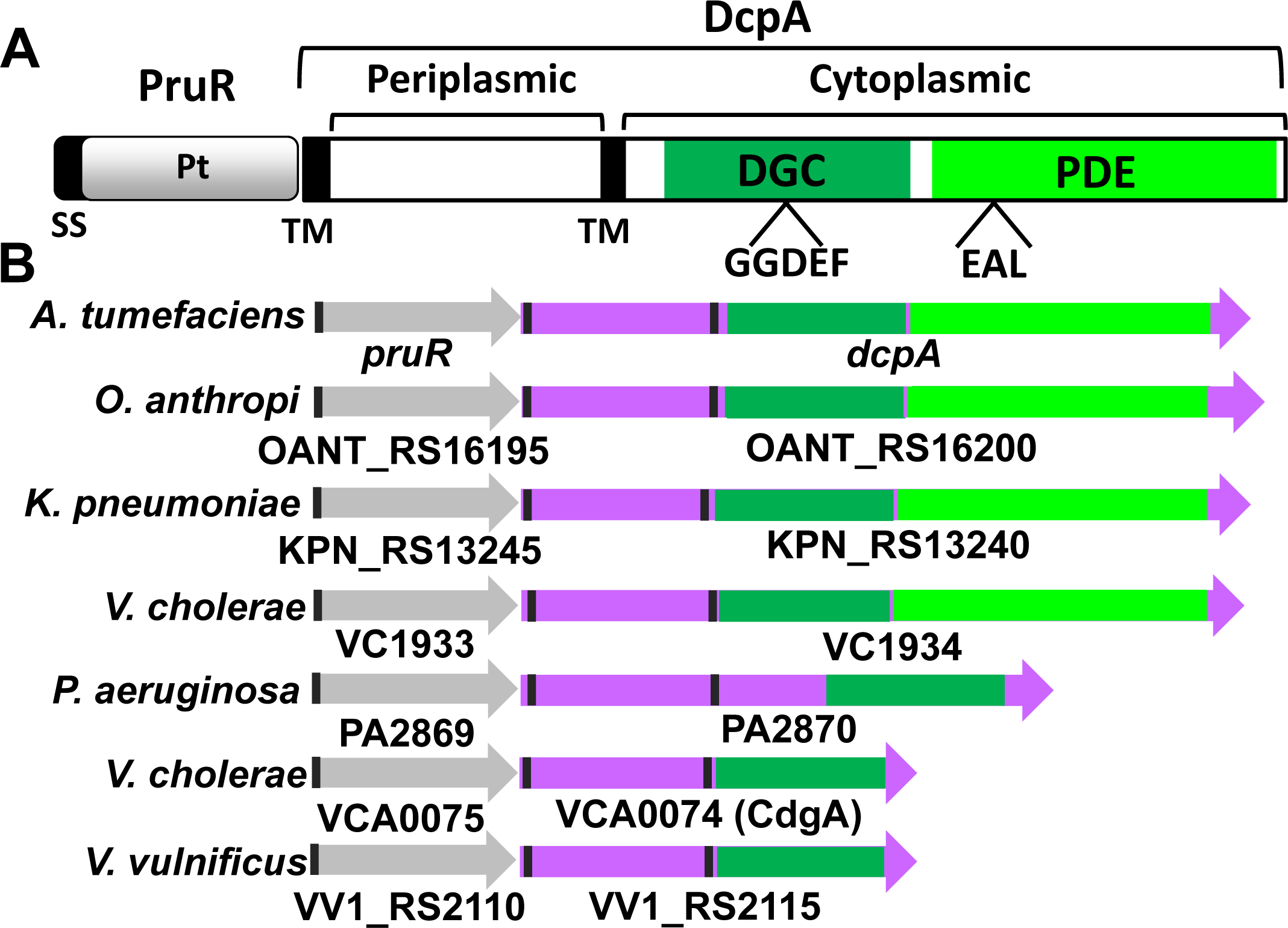
The *pruR-dcpA* operon is conserved across Proteobacteria in animal and human pathogens. **(A)** Domain structure for PruR and DcpA; transmembrane domains and signal sequence are black lines; DGC domain is green; PDE domain is lime green; Pt indicates the pterin-binding activity of the protein. **(B)** Presumptive operon structure for homologs of the *pruR-dcpA* operon from multiple pathogens. A subset of DcpA homologs are truncated relative to DcpA, and contain only the DGC domain. *Ochrobactrum anthropi* ATCC 49188, *Klebsiella pneumoniae pneumoniae* ATCC 700721; *Vibrio cholerae,* O1 biovar El Tor str. N16961*; Vibrio vulnificus* CMCP6; *Pseudomonas aeruginosa* PA01. Color coding as in panel A – gene and domain sizes are proportional.

### PruR-type proteins share a common structural fold

Based on our experimental findings for PruR-pterin interactions and on sequence conservation, we hypothesized that PruR represents a new class of proteins with an overall fold similar to the MoCo binding domains of SUOX-type proteins, that instead bind to the non-MoCo pterins. We purified representative members of the PruR family and determined their three-dimensional structures by X-ray crystallography (Table S2). The *A. tumefaciens* PruR structure reveals a general structural motif comprised of 10 β-sheets and 7 α-helices (Fig. 7A, PDB 7kou). The β-strands form one mixed, five-stranded β sheet (2-1-10-5-6) and one curved, four-stranded antiparallel β sheet (3-4-7-9), and these sheets are interconnected by short α helices (1, 2, 3, 5, 6 and 7). We solved an additional crystal structure of *A. tumefaciens* PruR (PDB 7kos) for which the overall structure is identical except that in this crystal form a small surface pocket is occupied by the side chain residues of neighboring or symmetry related polypeptide chains (Fig. S7 and Table S1).

Structures of three other PruR homologs were also solved including proteins from *V. vulnificus* (PDB 7kom, 30% identity; 1 Å), *V. cholerae* (PDB 7kp2, 32% identity, 1.03 Å), and *K. pneumoniae* (PDB 7rkb, 59% identity, 2.5 Å) (Fig. 6B and 6C). Multiple sequence and structure alignments of these PruR homologs clearly show that the structures are highly similar (root-mean-squared-deviation (RMSD) of 0.6 – 2.0 Å) and all share the SUOX-like fold. Overall, the PruR fold is conserved in different Alpha- and Gammaproteobacteria, consistent with a similar function for PruR in these bacteria.

**Figure 6.**
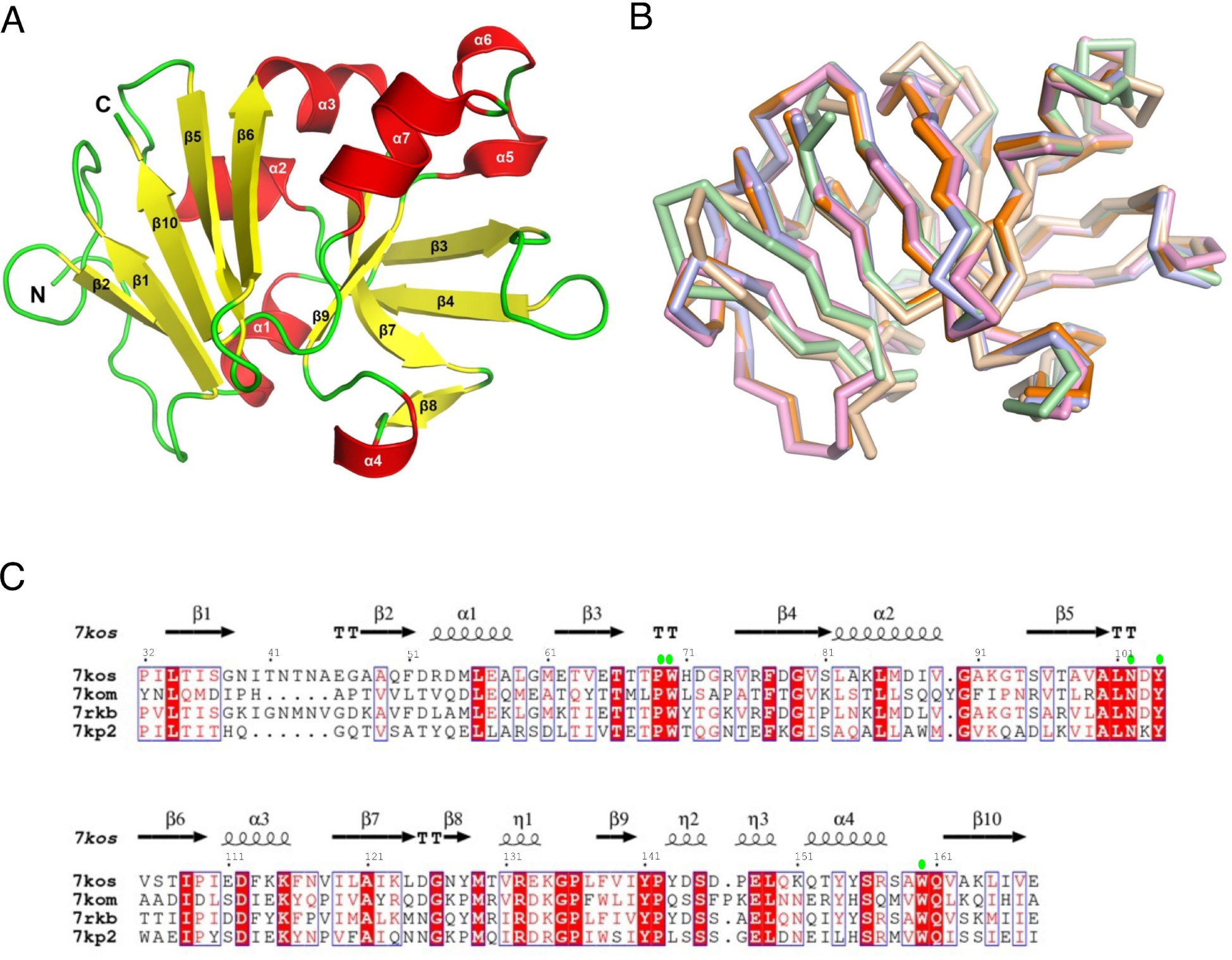
The structure of PruR is a degenerate SUOX fold conserved in multiple Alpha- and Gammaproteobacteria. **(A)** The overall structure of PruR from *A. tumefaciens* is depicted. The secondary structure elements are labeled and shown in red (α helices), yellow (β-strands) and green (loops). **(B)** Superposition of PruR from *A. tumefaciens* (orange, 7kos; violet, 7kou), *K. pneumoniae* (pink, 7rkb) *V. cholerae (*wheat, 7kp2), *V. vulnificus* (light green, 7kom). The peptide main chains of all structures are depicted as ribbons. **(C)** Multiple sequence alignment of PruR proteins with secondary structure elements from *A. tumefaciens* (PDB code 7kos) mapped above. Lime green circles mark pterin binding site residues conserved across all proteins.

**Figure 7.**
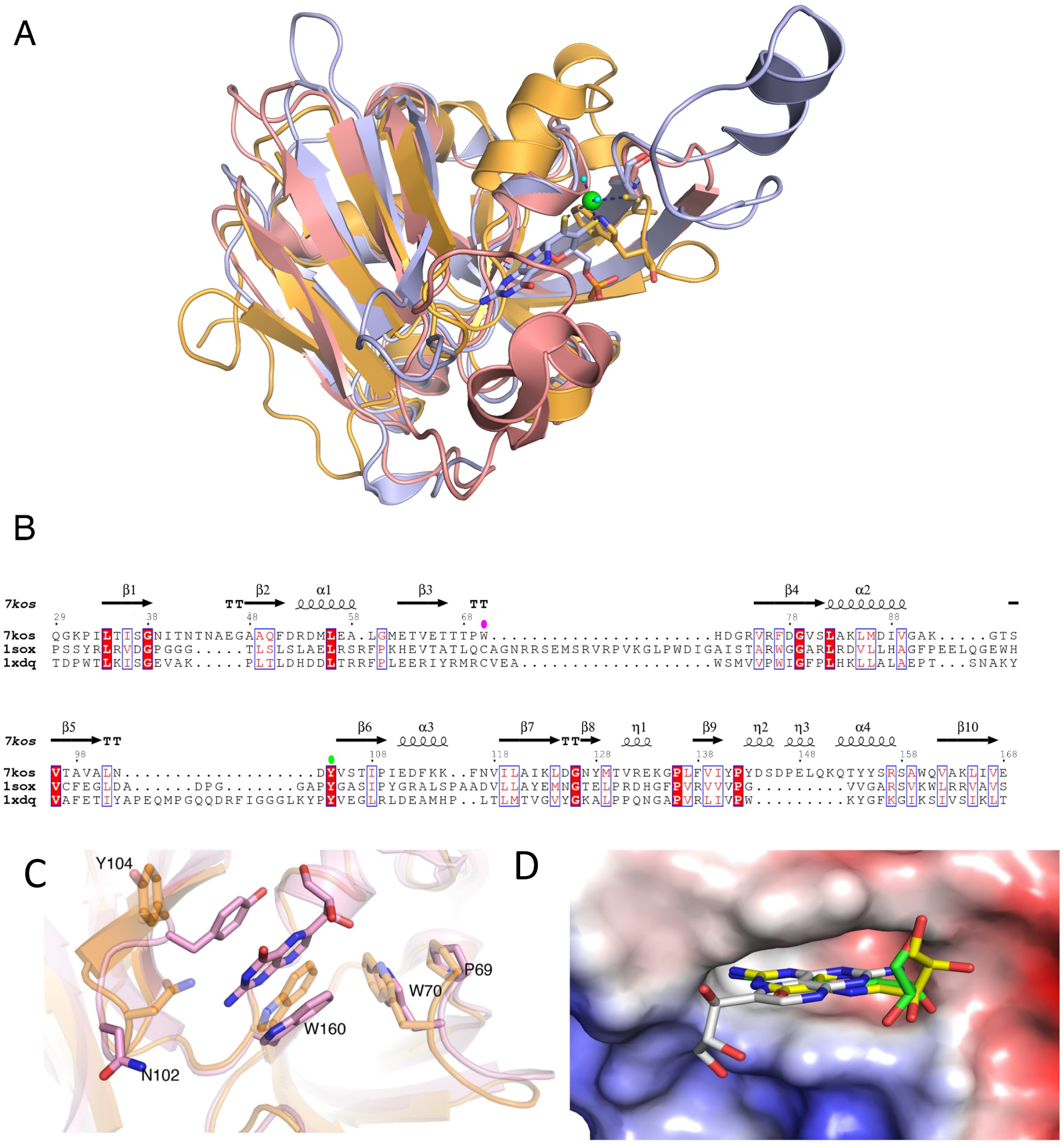
Comparison of PruR structure to conserved MoCo-binding domains. **(A)** Superposition of *A. tumefaciens* PruR (orange, 7kou) with the MoCo binding domain of chicken liver SUOX (violet, PDB 1sox) and *E. coli* YedY (salmon, PDB 1xdq). The MoCo and conserved C185 of SUOX are shown as balls-and-sticks (carbon, violet; oxygen, red; nitrogen, dark blue; sulfur, yellow), the molybdenum is shown as a green sphere, water molecules as small cyan spheres and hydrogen bonds as black, dashed lines. W70, conserved in PruR, is also shown as sticks. **(B)** Multiple sequence alignment of *A. tumefaciens* PruR with the MoCo binding domain of chicken liver sulfite oxidase and *E. coli* YedY. The magenta oval marks PruR W70 that aligns with the conserved **C185** in canonical MoCo-binding domains. The green oval marks an active site tyrosine conserved in pterin-binding proteins. **(C)** A zoomed in view of superposition of *A. tumefaciens* (orange, 7kou) and *K. pneumoniae* (pink, 7rkb) PruR. The neopterin and conserved residues of the pterin binding pocket are shown as sticks. Residue sidechains are numbered according to *A. tumefaciens*. **(D)** Superposition of all observed neopterin binding modes from crystal structures of *K. pneumoniae* (7RKB) and *V. cholerae* (7KP2). The binding pocket is represented as an electrostatic surface potential (blue-to-red, positive-to-negative charge) and neopterin as stick models. Carbons are in yellow (alternative conformation A, *V. cholerae*), grey (alternative conformation B, *V. cholerae*, 7kp2) and green (*K. pnuemoniae*, 7rkb), with oxygen in red and nitrogen in blue.

### PruR has a shortened MoCo binding site that associates with a pterin

Structural comparison of *A. tumefaciens* PruR to chicken liver sulfite oxidase (PDB 1sox), a relatively close SUOX structural homolog, and *E. coli* YedY (PDB 1xdq), a more distant sequence homolog, revealed significant structural overlap to the MoCo binding domains (Fig. 7A and 7B). PruR from *A. tumefaciens* aligns over 126 aa residues with the chicken liver SUOX protein with an RMSD value of 2.5 Å. The chicken liver SUOX active site contains a single molybdopterin bound to a surface cleft characterized by the conserved cysteine residue at the end of a lengthy β-hairpin (Fig. 7A). There is no surface cleft that could accommodate MoCo in the equivalent region of PruR, and the β-hairpin with the conserved Cys residue in all SUOX proteins, including in *E. coli* YedY (res. 138-146; (25), is truncated in PruR (res. 161-166). Furthermore, structural-sequence alignments of PruR, YedY, and chicken liver sulfite oxidase show that the canonical molybdenum-coordinating cysteine is substituted with a tryptophan (W70) in PruR, which constitutes a proline-tryptophan (^69^PW^70^) motif conserved in PruR homologs (Fig. 6C). Thus, the PruR fold is like that of SUOX family MoCo-binding domains but several structural determinants for MoCo binding, including a deep binding cleft, are lacking. The smaller pocket identified in our structures, however, can accommodate a smaller non-MoCo pterin molecule.

To examine the pterin-PruR interaction, we co-crystallized PruR with H_2_NPt, a commercially available monapterin stereoisomer that is more stable to oxidation than the tetrahydro pterins and thus compatible with crystallization methods. We obtained the ligand-bound structures for both *K. pneumoniae* and *V. cholerae* PruR homologs (7rkb and 7kp2, respectively) (Fig. 7C). Refinement revealed both were bound by the bicyclic moiety of the neopterin, with the hydroxylated tail absent. In all structures, the neopterin ring is positioned in the pterin binding cleft and is sandwiched between a tyrosine and tryptophan (*A. tumefaciens* Y104 and W160, respectively) (Fig. 7C). The pterin also forms hydrogen bonds with an asparagine residue (N102). These three residues are conserved across all PruR homologs (Fig. 6C), and Y104 is conserved in other SUOX proteins including YedY (Fig. 7B). Notably, in our apo structure of *A. tumefaciens* PruR, the Y104 side chain is flipped away from the pterin binding pocket (Fig. 7C), suggesting that binding of a ligand to this cleft induces key stacking interactions that stabilize the pterin in the site. Further comparison of the pterin-bound PruR structures also revealed three alternative binding conformations of neopterin. The orientation of the ring moiety varies, yielding different positions of the hydroxylated tail (Fig. 7D and Fig. S8A-C). We suspect that the observed variation in binding modes of neopterin is a result of differences between this disfavored fully oxidized ligand and the cognate reduced H_4_MPt and/or 2′-*O*-Met-H_4_MPt (9).

## DISCUSSION

Here we describe the regulation of the dual-function DGC/PDE DcpA by a novel pterin-binding protein PruR, building from our previous findings (9–11). All three components of the pterin regulatory pathway discussed here (PruR, DcpA, and PruA) were identified in a transposon mutagenesis screen designed to identify novel regulators of surface attachment in *A. tumefaciens* (10). In this study we have further interrogated this pterin regulatory pathway by (i) characterizing how the pterin-binding protein PruR regulates DcpA activity, (ii) defining PruR as a periplasmic protein that interacts with DcpA as well as H_4_MPt, (iii) elucidating the role of PruA in promoting the interaction between PruR and DcpA, and (iv) identifying similar regulatory systems in other *Proteobacteria*. Furthermore, we determined the three-dimensional structure of PruR and selected homologs in their apo forms, and two as a complex with a pterin. The level of conservation of PruR and DcpA homologs among multiple *Proteobacteria* suggests that they are likely to also utilize pterin-dependent regulation of cdGMP pools.

### A new class of pterin binding proteins

Our findings suggest that PruR and its homologs are a new type of pterin-binding periplasmic protein and we report the first structural information for this new group. The overall fold of PruR-type proteins suggests that they are a branch of the SUOX protein family, cytoplasmic enzymes including sulfite oxidases and nitrate reductases that utilize MoCo as a co-factor (16). Most SUOX proteins are large and have additional domains that drive catalysis. In contrast PruR-like proteins are small, almost entirely comprised of the SUOX fold and their N-terminal secretion signal. Pterins such as H_4_MPt and related molecules are significantly smaller and less structurally complex than MoCo. It is intriguing that the pterin binding site for PruR-type proteins and the SUOX MoCo binding site are structurally and likely evolutionarily related, especially given the markedly different functions of the two groups of proteins.

Other proteins known to bind the bicyclic pteridine ring are the aromatic amino acid hydroxylases such as phenylalanine hydroxylase (PAH), which coordinates a tetrahydrobiopterin (H_4_BPt) molecule as a cofactor. The H_4_BPt binding site stabilizes the pteridine ring via a conserved acidic residue and a histidine-conjugated iron atom, which interact with ring positions 2 and 4, respectively (26). This binding site thus bears no significant similarity to the SUOX-type pterin binding site of the PruR-type proteins we define here. There are also well studied, membrane-anchored mammalian proteins that associate with the essential metabolite tetrahydrofolate (H_4_F), which also contains the bicyclic pteridine ring. These folate binding proteins (FBPs) are upregulated in fetal cells and certain cancers (27, 28), and dramatically alter their conformation upon binding to H_4_F. Similar to PruR-like proteins, their association with the pteridine ring in H_4_F is fostered by conserved Tyr and Trp residues that sandwich the ring in a hydrophobic pocket, although these proteins are otherwise structurally distinct the PruR-type proteins reported here.

### Preference of PruR for fully reduced monapterin

The pterin binding specificity of PruR was revealed by *in vitro* studies with purified PruR and different pterin species. Fully reduced tetrahydropterins oxidize readily to the dihydropterin forms (29). We used a coupled enzyme reaction with the PruA pteridine reductase and NADPH in the presence of H_2_MPt and H_2_NPt under anoxic conditions to generate the respective tetrahydropterin species in the presence of PruR (11). We observed significantly greater association of the monapterin species in the presence of PruA than in its absence, consistent with greater binding to the PruA-generated H_4_MPt. In contrast, neopterin binding to PruR was lower and unaffected by the presence of PruA, even though our prior *in vitro* PruA work revealed similar catalytic efficiency with H_2_MPt and H_2_NPt substrates (11). PruR detectably binds the neopterin species and the reduced folate species, but not to the same extent as the PruA-generated H_4_MPt. Although this experiment is evidence for the pterin association with PruR, it does not provide binding affinities.

Co-crystallization with the H_2_NPt ligand and the PruR homologs from *V. cholerae* (7KOU) and *K. pneumoniae* (7RKB) revealed the fully oxidized neopterin (presumably due to oxidation of the exogenously added H_2_NPt during crystallization) binding at the analogous location as the MoCo binding site on SUOX. Pterin binding occurs within the pterin-binding cleft we have defined on the PruR-type proteins through interaction of the pteridine ring sandwiched between the conserved Tyr and Trp residues and several other hydrogen bonding positions. The hydroxylated tail of the neopterin was not visible in these structures suggesting greater mobility. The stereochemistry of the hydroxylated tail distinguishes monapterin from its stereoisomer neopterin, and the more complex benzoyl and glutamyl side chain substituents of folate, and we hypothesize that interactions between this tail and PruR add binding specificity. Additionally, the binding site may also interact directly with the fully reduced tetrahydropterin species through interactions that are not observed in the crystal structures with oxidized neopterin. The multiple positions of the neopterin detected within the binding cleft may also reflect weaker interactions of the protein with the non-cognate, fully oxidized neopterin molecule.

### PruR interacts with pterins in the periplasm

PruR negatively regulates surface attachment in *A. tumefaciens* through modulation of DcpA activity. Loss of function mutations in the pteridine reductase gene *pruA*, preventing H_4_MPt synthesis largely phenocopy *pruR* null mutants. The best studied functions for bacterial pterins are to act as enzymatic cofactors for amino acid hydroxylase enzymes such as phenylalanine hydroxylase (15). All evidence suggests that PruA drives reduction of H_2_MPt in the cytoplasm, but our findings have revealed that PruR functions and responds to pterins in the periplasm. A longstanding, unexplained observation among reports on bacterial pterins is that they accumulate extracellularly, with as much as 100-fold greater pterin in the extracellular fraction as compared to associated with cells (15, 30). The mechanism of excretion has not been elucidated, but it is also thought that extracellular pterins can re-enter cells (31). The PruR protein clearly interacts with a self-produced, reduced monapterin species in the periplasm, but exogenous pterins that enter the periplasm might also act as agonists or antagonists of the H_4_MPt-PruR interaction. Our findings are most consistent with H_4_MPt preferentially binding to PruR. However, H_4_MPt is susceptible to oxidation and thus will have a relatively short half-life in the periplasm and elsewhere outside the cell (32).

### Interaction of PruR and DcpA is stimulated by PruA

Crosslinking in whole cells reveals that PruR and DcpA interact through the DcpA periplasmic domain to form a stable complex. The size of the crosslinked species (∼85 kDa) suggests that the proteins are in a one-to-one stoichiometry. The crosslinked product between ectopically expressed PruR and the DcpA periplasmic domain is significantly diminished in the Δ*pruA* mutant, although it is still detectable. The Δ*pruA* mutant has a dramatic phenotype in which DcpA exhibits predominantly DGC activity, equivalent to the Δ*pruR* mutant (9), so whatever remaining interaction occurs between the two proteins is insufficient to maintain a strong DcpA PDE bias. Our analysis of the PruR control of DcpA catalytic site mutants suggests that in cells with active PruA, the PruR-DcpA interaction stimulates PDE activity and also diminishes DGC activity. We speculate this may be due the reciprocal control of dimerization for the separate DGC and PDE domains of DcpA, known to be required for these activities in other proteins that synthesize and degrade cdGMP (33, 34).

The DcpA periplasmic domain is less well conserved among the DcpA orthologs, although there are two invariant residues, Trp40 and Glu48 (in *A. tumefaciens* DcpA). The domains are all roughly 140 aa in length with dispersed chemical conservation. Alpha Fold predictions suggests that the DcpA periplasmic domain forms a novel four helix CACHE-type bundle, with overall structural similarity to the four helix bundles in the periplasmic domains of methyl-accepting dependent chemotaxis proteins (MCPs), that impart chemotactic motility responses to certain solutes (35). Rather than interacting directly with their ligands in the periplasm, several MCPs mediate the response through interactions with periplasmic solute binding proteins (SBPs) (36). The interaction of PruR and the DcpA periplasmic domain may share similarities with MCP-SBP interactions, with binding of the pterin fostering this interaction.

The PruR-DcpA system is strikingly analogous to the dual-function DGC-PDE MbaA in *V. cholerae*, which is regulated through the interaction of its periplasmic domain with NspS (37, 38). NspS is a SBP for polyamines and is encoded upstream of *mbaA* in the same operon. The specific type of polyamine bound by NspS can regulate its interaction with MbaA, thereby controlling the balance of DGC and PDE activities from this enzyme. *V. cholerae* synthesizes and releases norspermidine, which stimulates DGC activity through NspS-MbaA interactions, whereas other polyamines such as spermidine inhibit association with MbaA. (39, 40). The polyamine response impacts cdGMP levels, controlling *V. cholerae* biofilm formation. Interestingly, in the absence of polyamines, NspS can bind to MbaA and weakly impacts its activity, similar to our observation here that PruR weakly binds to DcpA in the absence of its proposed H_4_MPt ligand. There are multiple types of pterins produced and excreted by bacterial and eukaryotic organisms (12), and it is conceivable that similar to the NspS-MbaA system, the specific pterin-associated with PruR can impact its regulation of DcpA. These structurally variant pterins may act as agonists or antagonists for the PruR-H_4_MPt interaction. It is also plausible that the PruR-associated pterin acts as a proxy to detect a different cue in the periplasm, such as specific redox-active compounds.

### Insights into conserved PruR-DcpA type systems in other bacteria

The PruR homologs we identify here are well conserved proteins and highly likely to be orthologous in function. The structural determination of the four different PruR-type apo-proteins all had high similarity with an RMSD value of <2.0 Å. This similarity includes the residues we have identified that associate with the pteridine ring. Most of these *pruR* homologs are immediately upstream of *dcpA* homologs with both the DGC and PDE domains similar to DcpA, or only the DGC domain. The periplasmic domains of these DcpA-type proteins are of similar size and amino acid chemistry but with only the two fully conserved residues mentioned above. For some of the conserved *pruR-dcpA* operons, only one of the cdGMP functions is retained. For example, there are two of these conserved operons in *V. cholerae*, VC1933-VC1934 and VCA0075-VCA0074 (VCA0074 is the well-studied DGC known as CdgA, (24)). Interestingly VC1934 is unlikely to be active for cdGMP synthesis (due to its nonfunctional GADEF motif) but active for its degradation, and CdgA is well established to be active for cdGMP synthesis (24, 41). In combination they may provide similar, but independently regulated DGC/PDE activities, as does the dual functionality of DcpA in *A. tumefaciens.* In the two major model strains of *P. aeruginosa*, PA01 and PA14, mutants for their *dcpA* homologs (PA2780/PA1426970), both impact cdGMP-dependent phenotypes, although to different extents (42, 43). In all of these cases, the positioning of a *pruR*-type gene immediately upstream is consistent with a regulatory relationship with the downstream gene. Given the remarkable similarity of the PruR gene products, it is highly likely that they bind the same or related pterin ligands and provide pterin-dependent control.

## MATERIALS AND METHODS

Detailed descriptions of materials and methods can be found in the Supplementary Information, including: bacterial strains and plasmids used (Tables S3 and S4), growth conditions, molecular cloning and site-directed mutagenesis, Congo Red staining, biofilm formation quantification, PhoA secretion assays, antibody production and purification, western blots, periplasmic fractionation, *in vivo* crosslinking, HPLC analysis of pterins and folates, *in vitro* analysis of pterin binding to PruR, protein crystallography (including molecular cloning, protein production and purification, data collection, structure solution and refinement), purification of PruA, and *in vitro* enzymatic assays for DcpA.

## DATA AVAILABILITY

The structures determined here were deposited in the Protein Data Bank (https://www.rcsb.org/) with the assigned PDB codes: 7kos (*A. tumefaciens*), 7kou (*A. tumefaciens 2)*, 7kom (*V. vulnificus*), 7rkb (*K. pneumoniae*), 7kp2 (*V. cholerae*). Protein diffraction data have been deposited at proteindiffraction.org. All additional experimental data is available upon request from the authors.

## ACKNOWLEDGEMENTS

We thank Fitnat Yildiz, Seth Rubin, Vanessa Mariscal, Thomas Bernhardt, and Jon Beckwith for helpful discussions and Ramya Natarajan for technical assistance with preliminary protein purification. This project was supported by grants to C.F. (NIH R01 GM120337), K.D.A (NSF-CHE 2105598), A.K.G. (AI150466.). N.F. received support from the IU Genetics, Cellular and Molecular Sciences NIH Training Grant (T32 GM007757). Work by the Center for Structural Biology of Infectious Diseases was supported by HHS/NIH/NIAID contracts #HHSN272201700060C and 75N93022C00035. This research used resources of the Advanced Photon Source, a U.S. Department of Energy (DOE) Office of Science User Facility operated for the DOE Office of Science by Argonne National Laboratory under Contract No. DE-AC02-06CH11357. Use of the LS-CAT Sector 21 was supported by the Michigan Economic Development Corporation and the Michigan Technology Tri-Corridor (Grant 085P1000817). Access to LS-CAT and computational resources is facilitated by the NU Structural Biology Facility, which is supported by NCI grant to P30 CA060553 the Robert H. Lurie Comprehensive Cancer Center.

## REFERENCES

1. O. Ciofu, C. Moser, P. O. Jensen, N. Hoiby, Tolerance and resistance of microbial biofilms. Nat Rev Microbiol 20, 621–635 (2022).

2. K. Sauer et al., The biofilm life cycle: expanding the conceptual model of biofilm formation. Nat Rev Microbiol 20, 608–620 (2022).

3. C. Berne, C. K. Ellison, A. Ducret, Y. V. Brun, Bacterial adhesion at the single-cell level. Nat Rev Microbiol 16, 616–627 (2018).

4. U. Jenal, A. Reinders, C. Lori, Cyclic di-GMP: second messenger extraordinaire. Nat Rev Microbiol 15, 271–284 (2017).

5. T. E. Randall et al., Sensory perception in bacterial cyclic diguanylate signal transduction. J Bacteriol 204, e0043321 (2022).

6. M. A. Thompson, M. C. Onyeziri, C. Fuqua, Function and regulation of *Agrobacterium tumefaciens* cell surface structures that promote attachment. Curr Top Microbiol Immunol 418, 143–184 (2018).

7. S. B. Gelvin, *Agrobacterium*-mediated plant transformation: the biology behind the “gene-jockeying” tool. Microbiol. Mol. Biol. Rev. 67, 16–37 (2003).

8. J. E. Heindl et al., Mechanisms and regulation of surface interactions and biofilm formation in *Agrobacterium*. Frontiers in plant science 5, 176 (2014).

9. N. Feirer et al., A pterin-dependent signaling pathway regulates a dual-function diguanylate cyclase-phosphodiesterase controlling surface attachment in *Agrobacterium tumefaciens*. mBio 6, e00156 (2015).

10. J. Xu et al., Genetic analysis of *Agrobacterium tumefaciens* unipolar polysaccharide production reveals complex integrated control of the motile-to-sessile switch. Mol Microbiol 89, 929–948 (2013).

11. M. Labine et al., Enzymatic and mutational analysis of the PruA pteridine reductase required for pterin-dependent control of biofilm formation in *Agrobacterium tumefaciens*. J Bacteriol 10.1128/JB.00098-20 (2020).

12. N. Feirer, C. Fuqua, Pterin function in bacteria. Pteridines 28, 23–36 (2017).

13. J. M. Green, B. P. Nichols, R. G. Matthews, “Folate biosynthesis, reduction and polyglutamylation” in Escherichia coli and Salmonella: Cellular and Molecular Biology, F. C. Neidhardt, Ed. (ASM Press, Washington, D.C., 1996), vol. 1, chap. 41, pp. 665–673.

14. S. Leimkuhler, M. M. Wuebbens, K. V. Rajagopalan, The history of the discovery of the molybdenum cofactor and novel aspects of its biosynthesis in bacteria. Coord Chem Rev 255, 1129–1144 (2011).

15. A. Pribat et al., FolX and FolM are essential for tetrahydromonapterin synthesis in *Escherichia coli* and *Pseudomonas aeruginosa*. J Bacteriol 192, 475–482 (2010).

16. G. J. Workun, K. Moquin, R. A. Rothery, J. H. Weiner, Evolutionary persistence of the molybdopyranopterin-containing sulfite oxidase protein fold. Microbiol. Mol. Biol. Rev. 72, 228–248 (2008).

17. C. Kisker et al., A structural comparison of molybdenum cofactor-containing enzymes. FEMS Microbiol Rev 22, 503–521 (1998).

18. V. R. Sokya, W. Pfleiderer, R. Prewo, Pteridines: Synthesis and characteristics of 5,6-dihydro-6-(1,2,3-trihydroxypropyl)pteridines: Covalent intramolecular adducts. Helv. Chim. Acta. 73, 808–826 (1990).

19. C. Manoil, J. J. Mekalanos, J. Beckwith, Alkaline phosphatase fusions: sensors of subcellular location. J Bacteriol 172, 515–518 (1990).

20. H. Nielsen, K. D. Tsirigos, S. Brunak, G. von Heijne, A brief history of protein sorting prediction. Protein J 38, 200–216 (2019).

21. S. Okuda, H. Tokuda, Lipoprotein sorting in bacteria. Annu Rev Microbiol 65, 239–259 (2011).

22. D. Huber et al., Use of thioredoxin as a reporter to identify a subset of Escherichia coli signal sequences that promote signal recognition particle-dependent translocation. J Bacteriol 187, 2983–2991 (2005).

23. C. F. Schierle et al., The DsbA signal sequence directs efficient, cotranslational export of passenger proteins to the *Escherichia coli* periplasm via the signal recognition particle pathway. J Bacteriol 185, 5706–5713 (2003).

24. S. Beyhan, L. S. Odell, F. H. Yildiz, Identification and characterization of cyclic diguanylate signaling systems controlling rugosity in *Vibrio cholerae*. J Bacteriol 190, 7392–7405 (2008).

25. L. Loschi et al., Structural and biochemical identification of a novel bacterial oxidoreductase. J Biol Chem 279, 50391–50400 (2004).

26. J. A. Ronau et al., A conserved acidic residue in phenylalanine hydroxylase contributes to cofactor affinity and catalysis. Biochemistry 53, 6834–6848 (2014).

27. C. Chen et al., Structural basis for molecular recognition of folic acid by folate receptors. Nature 500, 486–489 (2013).

28. A. S. Wibowo et al., Structures of human folate receptors reveal biological trafficking states and diversity in folate and antifolate recognition. Proc Natl Acad Sci U S A 110, 15180–15188 (2013).

29. R. L. Blakley, S. J. Benkovic, Folates and Pterins: Chemistry and Biochemistry of Pterins (John Wiley & Sons, Inc., 1985), vol. 2.

30. K. Iwai, M. Kobashi, H. Fujisawa, Occurrence of *Crithidia* factors and folic acid in various bacteria. J Bacteriol 104, 197–201 (1970).

31. A. Noiriel, V. Naponelli, G. G. Bozzo, J. F. Gregory, 3rd, A. D. Hanson, Folate salvage in plants: pterin aldehyde reduction is mediated by multiple non-specific aldehyde reductases. Plant J 51, 378–389 (2007).

32. T. Arai et al., Auto-oxidation of 5,6,7,8-tetrahydroneopterin Pteridines 9, 26–28 (1988).

33. C. Chan et al., Structural basis of activity and allosteric control of diguanylate cyclase. Proc Natl Acad Sci U S A 101, 17084–17089 (2004).

34. F. Rao, Y. Yang, Y. Qi, Z. X. Liang, Catalytic mechanism of cyclic di-GMP-specific phosphodiesterase: a study of the EAL domain-containing RocR from *Pseudomonas aeruginosa*. J Bacteriol 190, 3622–3631 (2008).

35. A. Ortega, I. B. Zhulin, T. Krell, Sensory repertoire of bacterial chemoreceptors. Microbiol Mol Biol Rev 81 (2017).

36. J. P. Cerna-Vargas, B. Sanchez-Romera, M. A. Matilla, A. Ortega, T. Krell, Sensing preferences for prokaryotic solute binding protein families. Microb Biotechnol 16, 1823–1833 (2023).

37. A. A. Bridges, J. A. Prentice, C. Fei, N. S. Wingreen, B. L. Bassler, Quantitative input-output dynamics of a c-di-GMP signal transduction cascade in *Vibrio cholerae*. PLoS Biol 20, e3001585 (2022).

38. E. Karatan, T. R. Duncan, P. I. Watnick, NspS, a predicted polyamine sensor, mediates activation of *Vibrio cholerae* biofilm formation by norspermidine. J Bacteriol 187, 7434–7443 (2005).

39. A. A. Bridges, B. L. Bassler, Inverse regulation of *Vibrio cholerae* biofilm dispersal by polyamine signals. Elife 10 (2021).

40. S. R. Cockerell et al., *Vibrio cholerae* NspS, a homologue of ABC-type periplasmic solute binding proteins, facilitates transduction of polyamine signals independent of their transport. Microbiology 160, 832–843 (2014).

41. B. Lim, S. Beyhan, F. H. Yildiz, Regulation of Vibrio polysaccharide synthesis and virulence factor production by CdgC, a GGDEF-EAL domain protein, in Vibrio cholerae. J Bacteriol 189, 717–729 (2007).

42. D. G. Ha, M. E. Richman, G. A. O’Toole, Deletion mutant library for investigation of functional outputs of cyclic diguanylate metabolism in *Pseudomonas aeruginosa* PA14. Appl Environ Microbiol 80, 3384–3393 (2014).

43. K. Eilers et al., Phenotypic and integrated analysis of a comprehensive *Pseudomonas aeruginosa* PAO1 library of mutants lacking cyclic-di-GMP-related genes. Front Microbiol 13, 949597 (2022).

