## Supplemental Files and Figures for "Control of Biofilm Formation by an *Agrobacterium tumefaciens* Pterin-Binding Periplasmic Protein Conserved Among Pathogenic Bacteria"

Running Title: Pterin-Dependent Regulation of Biofilm Formation

Supplementary Tables: 4
Supplementary Methods
Supplementary References
Supplementary Figure Legends
Supplementary Figures: 8

#### Supplementary Tables

#### Table 1. Crystallographic and structural information for *A. tumefaciens* PruR

| **PDB Accession Code** | **7kos – *A. tumefaciens*** | **7kou – *A. tumefaciens* (structure variant)** |
|---|---|---|
| **Data Collection** | | |
| Space group | $P2_12_12_1$ | $P2_12_12$ |
| Unit cell parameters (Å; º) | $a = 80.46, b = 91.81, c = 99.22;$ | $a = 120.93, b = 62.02, c = 101.92;$ |
| | $\alpha = 90.00, \beta = 90.00, \gamma = 90.00$ | $\alpha = 90.00, \beta = 90.00, \gamma = 90.00$ |
| Resolution range (Å) | 30.00 - 1.50 (1.53 - 1.50) | 30.00 - 1.83 (1.86 - 1.83) |
| No. of reflections | 118,548 (5,876) | 68,046 (3,378) |
| $R_{\text{merge}}$ (%) | 4.6 (79.6) | 8.4 (77.3) |
| Completeness (%) | 99.9 (100.0) | 100.0 (100.0) |
| $\langle I/\sigma(I)\rangle$ | 31.3 (2.4) | 19.4 (2.2) |
| Multiplicity | 5.8 (5.8) | 5.4 (5.4) |
| Wilson $B$ factor | 21.7 | 20.4 |
| **Refinement** | | |
| Resolution range (Å) | 29.04 - 1.50 (1.54 - 1.50) | 29.80 - 1.83 (1.88 - 1.83) |
| Completeness (%) | 99.9 (100.0) | 99.7 (96.5) |
| No. of reflections | 111,854 (8,594) | 64,577 (4,806) |
| $R_{\text{work}}/R_{\text{free}}$, (%) | 15.2/17.2 (26.9/27.9) | 16.2/19.3 (26.9/28.7) |
| Protein chains/atoms | 4/4,581 | 4/4,546 |
| Ligand/Solvent atoms | 88/736 | 45/735 |
| Mean temperature factor (Å$^2$) | 25.2 | 23.7 |
| **Coordinate Deviations** | | |
| R.m.s.d. bonds (Å) | 0.006 | 0.006 |
| R.m.s.d. angles (º) | 1.435 | 1.364 |

| Ramachandran plot | | |
|---|---|---|
| Favored (%) | 98.0 | 98.0 |
| Allowed (%) | 2.0 | 2.0 |
| Outside allowed (%) | 0.0 | 0.0 |

#### Table 2. Crystallographic and structural information for PruR homologs

| PDB Accession Code | 7kom – *V. vulnificus* | 7rkb – *K. pneumoniae* | 7kp2 – *V. cholerae* |
|---|---|---|---|
| **Data Collection** | | | |
| Space group | *I222* | $P4_12_12$ | $P2_1$ |
| Unit cell parameters (Å; º) | *a = 33.05, b = 71.09, c = 106.36;* | *a = 83.52, b = 83.52, c = 45.88;* | *a = 33.49, b = 54.62, c = 33.45;* |
| | $\alpha = 90.00, \beta = 90.00, \gamma = 90.00$ | $\alpha = 90.00, \beta = 90.00, \gamma = 90.00$ | $\alpha = 90.00, \beta = 91.5, \gamma = 90.00$ |
| Resolution range (Å) | 30.00 - 0.99 (1.01 - 0.99) | 30.00 - 2.50 (2.54 - 2.50) | 30.00 - 1.03 (1.05 - 1.03) |
| No. of reflections | 65,367 (2,840) | 5,990 (298) | 57,114 (2,778) |
| $R_{merge}$ (%) | 7.5 (67.9) | 15.7 (167.5) | 9.2 (76.4) |
| Completeness (%) | 93.4 (82.2) | 100.0 (100.0) | 96.5 (93.4) |
| $\langle I/\sigma(I) \rangle$ | 45.1 (3.2) | 20.1 (2.2) | 22.1 (3.2) |
| Multiplicity | 7.3 (6.0) | 13.7 (14.2) | 4.6 (4.3) |
| Wilson *B* factor | 6.8 | 57.0 | 7.9 |
| **Refinement** | | | |
| Resolution range (Å) | 26.60 - 0.99 (1.02 - 0.99) | 29.53 - 2.50 (2.57 - 2.50) | 21.48 - 1.03 (1.06 - 1.03) |
| Completeness (%) | 93.2 (82.9) | 99.7 (97.7) | 96.3 (91.0) |
| No. of reflections | 62,187 (4,270) | 5,653 (424) | 54,176 (3,973) |
| $R_{work}/R_{free}$, (%) | 12.0/14.1 (26.0/30.1) | 19.9/25.7 (27.3/41.0) | 13.1/14.8 (23.1/23.9) |
| Protein chains/atoms | 1/1,098 | 1/1,161 | 1/1,096 |
| Ligand/Solvent atoms | 7/254 | 37/14 | 36/189 |

| | | | |
|---|---|---|---|
| Mean temperature factor ($Å^2$) | 9.1 | 63.0 | 11.6 |
| **Coordinate Deviations** | | | |
| R.m.s.d. bonds (Å) | 0.007 | 0.003 | 0.007 |
| R.m.s.d. angles (º) | 1.53 | 1.182 | 1.486 |
| **Ramachandran plot** | | | |
| Favored (%) | 99.0 | 96.0 | 99.0 |
| Allowed (%) | 1.0 | 4.0 | 1.0 |
| Outside allowed (%) | 0.0 | 0.0 | 0.0 |

**Table S3: Strains and Plasmids**

| Strain/Plasmid | Relevant Information | Source |
|---|---|---|
| ***E. coli*** | | |
| DH5α/λpir | *λpir*; Cloning strain | (1) |
| BL21(DE3) (Magic) | Expression strain for T7-promoter expression system | (2) |
| BL21 Codon Plus (DE3)-RIL | Expression strain for T7-promoter expression system | Agilent |
| BL2-Gold (DE3) | Expression strain for T7-promoter expression system | Agilent |
| ***A. tumefaciens*** | | |
| C58 | Nopaline type strain; pTIC58; pAtC58 | (3) |
| NF003 | C58Δ*pruR* (polar on *dcpA*) | (4) |
| JX105 | C58Δ*cel*Δ*upp* | (5) |
| JX137 | C58Δ*pruA* | (5) |
| JX138 | C58Δ*dcpA* | (5) |
| JX165 | C58Δ*cel*Δ*upp*Δ*pruA* | This study |
| JE009 | C58Δ*pruR* (non-polar) | This study |
| JE014 | C58Δ*pruRdcpA* | This study |
| **Plasmids** | | |
| pET-15b | T7 expression plasmid for protein purification; $Ap^R$ | Novagen (EMD Millipore) |
| pSRKGm | Broad host range $P_{lac}$ expression vector; *lacIQ*; $Gm^R$ | (6) |
| pNPTS138 | ColE1 suicide vector; *sacB*; $Km^R$ | (7) |
| pMCSG53 | Vector with N-terminal $His_6$ and TEV protease; $Ap^R$ | (8) |
| pTYB12 | T7 expression plasmid for N-terminal intein tag fusions; $Ap^R$ | New England BioLabs |
| pMAL-c2 | $P_{tac}$ inducible plasmid for N-terminal maltose-binding protein fusions; *lacIQ*; $Ap^R$ | New England Biolabs |
| pNF034 | pET15b carrying $His_6$-*pruA*; $Ap^R$ | (4) |
| pNF036 | pSRKGm carrying $P_{lac}$-*pruR-dcpA*; $Gm^R$ | (4) |
| pNF038 | pSRKGm carrying $P_{lac}$-*pruR*; $Gm^R$ | (4) |
| pJX146 | pSRKGm carrying $P_{lac}$-*pruA*; $Gm^R$ | (5) |
| pNF030 | pSRKGm carrying *Plac-dcpA*$^{E308A}$; GmR | (4) |
| pNF031 | pSRKGm carrying *Plac-dcpA*$^{E431A}$; GmR | (4) |
| pNF032 | pSRKGm carrying $P_{lac}$-*dcpA*$^{E308A\ E431A}$; $Gm^R$ | (4) |
| pNF063 | pET15b carrying $His_6$-*pruR*$_{\Delta SS}$; $Ap^R$ | This study |
| pNF068 | pSRKGm carrying $P_{lac}$-*pruR-phoA*; $Gm^R$ | This study |
| pNF073 | pSRKGm carrying $P_{lac}$-*pruR*$_{SS}$-*phoA*; $Gm^R$ | This study |
| pNF084 | pSRKGm carrying $P_{lac}$-*dcpA*$^{cyt}$; $Gm^R$ | This study |
| pNF090 | pMAL-c2 carrying $P_{tac}$- *MBP-dcpA*$^{cyt}$; $Ap^R$ | This study |

| | | |
|---|---|---|
| pNF096 | pSRKGm carrying *$P_{lac}$-pruR$_{\Delta SS}$-dcpA*; $Gm^R$ | This study |
| pNF098 | pSRKGm carrying *$P_{lac}$-pruR$^{C19A}$-dcpA*; $Gm^R$ | This study |
| pNF108 | pTYB12 carrying *$P_{T7}$-dcpA$^{peri}$*; $Ap^R$ | This study |
| pJE005 | pNPTS138 carrying *pruR* SOE deletion fragment; $Km^R$ | This study |
| pJE007 | pSRKGm carrying *$P_{lac}$-dcpA$^{peri}$*; $Gm^R$ | This study |
| pCF682 | pSRKGm carrying *$P_{lac}$-malE$_{SS}$-pruR-3XFLAG*; $Gm^R$ | This study |
| pCF683 | pSRKGm carrying *$P_{lac}$-dsbA$_{SS}$-pruR-3XFLAG*; $Gm^R$ | This study |
| pMCSG53-PruR | pMCSG53 carrying residues 23-168 of *pruR* from *A. tumefaciens* | This study |
| pMCSG53-KpnPruR | pMCSG53 carrying residues 24-169 of putative *pruR* homologue from *Klebsiella pneumoniae subsp. Pneumoniae NTUH-K2044* | This study |
| pMCSG53-VchPruR | pMCSG53 carrying residues 22-156 of putative *pruR* homologue from *Vibrio cholerae O1 biovar El tor st. N16961* | This study |
| pMCSG53-VvulPruR | pMCSG53 carrying residues 19-153 of putative *pruR* homologue from *V. vulnificus CMCP6* | This study |

**Table S4: Oligonucleotides Used in This Study**

| **Primer** | **Sequence** |
|---|---|
| *pruA* deletion P1 | ACTAGTGCGCGATTATCCGCGGTAATC |
| *pruA* deletion P2 | AAGCTTGGTACCGAATTCTGAACGGAAAGAAGCTGCCTGATG |
| *pruA* deletion P3 | GAATTCGGTACCAAGCTTCATACACGCCCTATTTGCTGG |
| *pruA* deletion P4 | CTGCAGAGTCATAGTTGACCGTACCACCGA |
| *pruR* deletion2 P1 | ACGCCAAGCTACGTAATACGACTCACTAGTGCGGAACAGATGTCTGGATATTTCCG |
| *pruR* deletion2 P2 | CTTTATTCCACTACACTCATATTTTTTCTCCATCTATAACGTTCAG |
| *pruR* deletion2 P3 | AAAATATGAGTGTAGTGGAATAAAGGTGCCGTTTGCGAAA |
| *pruR* deletion2 P4 | GCCTTGACTAGAGGGTCGACGCATGCGGTATAGACGATATGCAGCTTGCGA |
| PruR-DcpA P1 | ATCATATGAGTGTCGGCCGGGTACTTC |
| PruR-PhoA 5' | GCAAGCTTGTGTTATTCCACTATCAGTTTGG |
| PruR-PhoA Whole 5' | GACTGCAGACTCGGACACCAGAAATGC |
| PruR-PhoA Whole 3' | GCTAAGCTTGTGTTATTTCAGCCCCAGAGC |
| PruR Upstream 5' | GATCATATGCGTTGAGTCACACCGGACGGATTGC |
| PruR Upstream 3' | GAACTGCAGTATGGCCTGCGCGCAGGAAAG |
| *dcpA$^{cyt}$* F | GCTGGAATTGCAGGCGAAG |
| *dcpA$^{cyt}$* R | GGTCTCCATCACCCATTTTC |
| *dcpA* pMAL F | GATGGATCCGAGCAGAACGATCTCCTGAACC |
| dcpA pMAL R | GCTGAGCTCGATCAGACCGAACTCCAAGCC |
| pruRdel$_{SS}$-dcpA F | CTCCATATGTGCGGGACAATTGCCAAAC |
| pruRdel$_{SS}$-dcpA R | GCGACTAGTGCTTATTCCACTATCAGTTTGG |

| | |
|---|---|
| dcpA$^{peri}$ F | TAACAATTTCACACAGGAAACAGCATATGTGCGAAAACTTGTGGCGGTGA |
| dcpA$^{peri}$ R | GAATTCCTGCAGCCCGGGGGATCCACTAGTTCAATCGTTCTGGCGCAGCA |
| *dcpA-peri-intein F* | GGAATTCGAGCGGCAGACCG |
| *dcpA-peri-intein R* | GTCCCGGGTATATGCAGCTTGCGAACATC |
| *pruR*$^{C19A}$ F | CTTATGTGGCTTTCCGCCGCGCAGGCCGGGAC |
| *pruR*$^{C19A}$ R | GTCCCGGCCTGCGCGGCGGAAAGCCACATAAG |
| *MalE$_{SS}$-pruR* F | ATTTCACACAGGAAACAGCATATGAAAATAAAAACAGGTGCACGCATC |
| *MalE_DsbA$_{SS}$-pruR* R | CCTGCAGCCCGGGGGATCCACTAGTTTATTCCACTATCAGTTTGGCAACTTGCCATGCGGA |
| *DsbA$_{SS}$-pruR* F | ATTTCACACAGGAAACAGCATATGAAAAAGATTTGGCTGG |
| *His6-pruRnoSS F* | GATGGATCCGCTCAGACCGAACTCCAAG |
| *His6-pruRnoSS R* | CTCCATATGTGCGGGACAATTGCCAAAC |
| pMCSG53-PruR F | TACTTCCAATCCAATGCCGACCAGGTCGCTTTCTTTCACG |
| pMCSG53-PruR R | TTATCCACTTCCAATGTTAGGCCAGTAGCGTAATCGGATGTT |
| pMCSG53-KpnPruR F | TACTTCCAATCCAATGCCGAAAAACTGCCTGCGCCGACA |
| pMCSG53-KpnPruR R | TTATCCACTTCCAATGTTATTCTATAATCATCTTTGAAACTTGCCAGGC |
| pMCSG53-VchPruR F | TACTTCCAATCCAATGCCCAAGAACCGATCCTCACGATAACC |
| pMCSG53-VchPruR R | TTATCCACTTCCAATGTTAAGGAGTGATGATTTCTATACTGCTGATCT |
| pMCSG53-VvulPruR F | TACTTCCAATCCAATGCCTATAATTTACAAATGGATATCCCCCATGCC |
| pMCSG53-VvulPruR R | TTATCCACTTCCAATGTTACTTCGCGATATGTATTTGTTTTAACTGCCAAA |

#### Supplemental Materials and Methods

**Strains, plasmids, and growth conditions.** All strains, plasmids, and oligonucleotides used in this study are listed in the supplemental material, in Table S3 and S4. Oligonucleotides were synthesized by Integrated DNA Technologies (Coralville, IA). All sequencing was done by the Indiana Molecular Biology Institute (Bloomington, IN) or ACGT (Wheeling, IL) as noted. *A. tumefaciens* strains were grown in AT minimal media (9) supplemented with 0.5% (w/v) glucose and 15 mM ammonium sulfate) at 30$^{o}$C.

**Cloning and site-directed mutagenesis**. Ectopic expression of *pruR* and *dcpA* was controlled by cloning the coding sequence into pSRKGm, which contains an inducible $P_{lac}$ promoter and *lacI$^{q}$* using the primers listed in Table S4 and the NdeI and SpeI restriction sites. Site directed mutagenesis of the codons for the catalytic residues of DcpA and the potential lipidation site of PruR was done using a QuikChangeII Site Directed Mutagenesis Kit (Agilent, La Jolla, CA) following manufacturer's instructions with the plasmid containing the wild-type *dcpA* or *pruR* allele as the template. Plasmids were then transformed into chemically competent *E. coli* DH5α and plated on Lenox Broth solid media (10 g/L tryptone, 5 g/L sodium chloride, 5 g/L yeast extract, 15 g/L agar) containing 50 µg/mL gentamicin. Following sequencing by the Indiana Molecular Biology Institute, plasmids were transformed via electroporation into *A. tumefaciens* and then plated onto ATGN solid medium (ATGN media supplemented with 1.5% agar) containing 300 µg/mL gentamicin.

To construct a markerless, in-frame deletion of *pruR*, the upstream region and the downstream region, including the last nine nucleotides of *pruR* were amplified and ligated into pNPTS138 digested with SphI and SpeI using the NEBuilder Hi-Fi DNA Assembly Kit (New England Biolabs, Ipswich, MA) following the manufacturer's protocol and transformed into chemically competent *E. coli* DH5α and plated on Lenox Broth solid media containing 50 μg/mL kanamycin. Following sequencing by ACGT (Wheeling, IL), the plasmid was introduced into *A. tumefaciens* via conjugation. Kanamycin-sensitive, sucrose resistant colonies were confirmed to have the mutation using primers *pruR* deletion2 P1 and *pruR* deletion2 P4 (Table S4).

**Congo Red Staining Assay.** 3 μL of early stationary phase cultures of *A. tumefaciens* were spotted onto ATGN plates supplemented with 75 μg/mL Congo Red with either no additions or 400 μM IPTG to induce plasmid expression. Plates were incubated for 48 h at 30 °C prior to photographing.

**Biofilm Formation Quantification.** Polyvinyl chloride coverslips were placed upright in a 12-well tissue culture plate (Corning, Inc.) and UV sterilized. Mid-exponential *A. tumefaciens* cultures were sub-cultured to a starting $OD_{600}$ of 0.05 in ATGN media supplemented with 22 μM iron sulfate and 400 μM IPTG to induce plasmid expression, then incubated statically at room temperature for 48 h in a humidity-controlled chamber. To quantify cellular attachment, coverslips were washed with water and stained with 0.1 % (w/v) crystal violet (CV). To measure adhered cells, the CV was solubilized in 1 mL of 33% acetic acid, and then the absorbance at 600 nm ($A_{600}$) was measured. The $A_{600}$ measurement of acetic acid was subtracted from readings prior to normalization against the $OD_{600}$ of planktonic cells remaining in the well, with the absorbance of the media subtracted out.

**PhoA secretion assays.** Alkaline phosphatase assays to evaluate protein secretion were based on a previously published procedure (10). Briefly, overnight *A. tumefaciens* cultures containing *phoA* fusion constructs were subcultured 1:10 into ATGN media with 400 µM isopropyl β-1-thiogalactopyranoside (IPTG) and incubated at 28 °C until mid-exponential phase. After measurement of culture $OD_{600}$, 2 mL of culture was centrifuged (7800 *g* x 3 min.) and resuspended in 2 mL 1 M Tris-HCl (pH 8.0) on ice. 400 µL of resuspended culture was then added to 600 µL of 1 M Tris-HCl (pH 8.0), for a total 1 mL reaction volume. Iodoacetamide was added to a concentration of 1 mM to prevent functional folding of cytoplasmic alkaline phosphatase in non-growing cells. After resuspension, one drop of 0.1% (w/v) SDS and two drops of chloroform were added and cell mixtures were vortexed for five seconds. After vortexing, 100 µL of 0.4% (w/v, in 1 M Tris-HCl, pH 8.0) para-nitrophenylphosphate (PNPP) was added, cultures vortexed again, and incubated at 37°C. After the development of light yellow color, reactions were stopped by the addition of 100 µL 1 M $KH_2PO_4$ and the total time of the reaction was recorded, and reaction absorbance at 420 and 550 nM was determined using a Biotek Synergy HT microplate reader. Alkaline phosphatase specific activity (in Miller units) was calculated as; $1000 * A_{420} / (OD_{600} * t * f)$, with *t* representing time in minutes and *f* representing the fractional volume of cell suspension/(volume of cells + reaction buffer) in the reaction.

**Protein purification of Intein-tagged proteins.** An overnight culture of BL-21(DE3) containing the corresponding pTYB12 derivative was subcultured 1:100 into 1 L LB containing 100 µg/mL ampicillin. Culture was grown at 37 °C to OD600: 0.5, at which time IPTG was added to a final concentration of 200 µM to induce protein expression. Culture was grown at 16°C for 16-18 h and collected via centrifugation (5,600 *g* x 15 min., 4°C). Culture pellets were weighed and stored at -80°C until the purification procedure was continued. Pellets were thawed, resuspended in 10 mL/g chitin column buffer (20 mM Tris-HCl, 200 mM NaCl, 1 mM EDTA, pH 8.5), PMSF added to a final concentration of 1 mM, and lysed using a M-110L Microfluidizer Processor (Microfluidics, Westwood, MA). Cell lysates were centrifuged (14,200 *g* x 1hr, 4°C) to remove cell debris and the clarified extract was collected. The clarified extract was diluted 1:4 with chitin column buffer and loaded (using gravity flow) onto a column containing chitin resin (New England BioLabs, Ipswich, MA) that had been previously equilibrated and washed with chitin column buffer. After loading, the column was washed with chitin column buffer and wash and flowthrough fractions collected to determine protein binding efficiency. Three bed volumes (~30 mL) of cleavage buffer (20 mM Tris-HCl, 200 mM NaCl, 1 mM EDTA, 50 mM DTT, pH 8.5) were run through the column, at which point flow was stopped. The column was then incubated with cleavage buffer at room temperature for 40 hours to allow self-cleavage of the intein tag to take place. After incubation, protein was eluted using chitin column buffer. 1.5 mL elution fractions were collected and separated by SDS-PAGE (5% stacking, 10% resolving) to determine fraction purity and yield. Fractions containing the protein of interest were pooled and dialyzed into chitin column buffer with 1 mM DTT using Slidea-Lyzer 3000 MWCO dialysis cassettes (ThermoFisher, Waltham, MA). Glycerol was added to a final concentration of 5%, samples were flash frozen, and stored at -80°C until use.

**Protein purification of maltose binding protein (MBP)-tagged DcpA$^{cyt}$.** An overnight culture of BL-21(DE3) containing the corresponding pMAL derivative was subcultured 1:100 into 1 L Lenox Broth containing 0.2% glucose plus 100 µg/mL ampicillin. Culture was grown at 37 °C to $OD_{600}$ of 0.5, at which time IPTG was added to a concentration of 400 µM to induce protein expression. Culture was grown at 16 °C for 16-18 h and collected via centrifugation (5,600 *g* x 15 min, 4°C). Culture pellets were weighed and stored at -80 °C until the purification procedure was continued. Pellets were thawed, resuspended in 10 mL/g buffer A (20 mM Tris-HCl (pH 7.4), 200 mM NaCl, 1 mM EDTA, and 1 mM DTT), PMSF added to a final concentration of 1 mM, and lysed using a M-110L Microfluidizer Processor (Microfluidics, Westwood, MA). Cell lysates were centrifuged (14,200 *g* x 1hr, 4°C) to remove cell debris and the clarified extract was collected. The clarified extract was diluted 1:6 with buffer A and loaded (using gravity flow) onto a column containing amylose resin (New England BioLabs, Ipswich, MA) that had been previously equilibrated and washed with Buffer A. After loading, the column was washed Buffer A and wash and flow-through fractions collected to determine protein binding efficiency. Protein was then eluted using Buffer A + 10 mM maltose. 2-3 mL elution fractions were collected separated by SDS-polyacrylamide (10% resolving, 5% stacking) to determine fraction purity and yield. Fractions containing the protein of interest were pooled and concentrated using Amicon Ultra 15 mL centrifugal filter units

(Millipore, Darmstadt, Germany). Glycerol was added to a final concentration of 5%, samples were flash frozen, and stored at -80°C until use.

**Size Exclusion Chromatography.** A Superdex 200 (DcpA) or 75 (PruR) 10/300 GL gel filtration column was utilized to determine the size and oligomerization state of purified proteins. Before loading protein samples, an ÄKTA FPLC system (GE Healthcare Bio-Sciences, Pittsburgh, PA) was used to equilibrate the column with two column volumes (~50 mL) of buffer (either TEDG- 50mM Tris-HCl pH 7.5, 0.2M NaCl, 5% glycerol, 1mM DTT, and 0.5mM EDTA or amylose column buffer- 20mM Tris pH=7.4, 0.2M NaCl, 1mM EDTA, and 1mM EDTA). After equilibration, 250 μL of 15-600 protein standard mix (Sigma-Aldrich, St. Louis, MO) was loaded onto the column. Two column volumes of buffer were applied at 0.5 mL/min to the column and protein elution was monitored via UV detection. The column was washed and then regenerated until no further UV signal was observed and re-equilibrated with the amylose column buffer. 250 μL of purified protein (specific concentrations in corresponding Fig. legend) was then loaded at 0.5 mL/min to the column. Two column volumes of buffer were applied to the column (0.5 mL/min), during which 250 μL elution fractions were collected and protein elution monitored via UV detection. To determine protein size, experimental peak elution times were calculated using a standard curve of elution times of the protein standards. Peak elution fractions were separated by SDS- PAGE and gels were stained with Coomassie Blue.

***In vitro*** **DGC and PDE enzymatic assays.** These assays were performed using modifications of the EnzChek Pyrophosphate Assay (Thermo Fisher, Waltham, MA).

This system measures the presence of inorganic phosphate using purine nucleotide phosphorylase (PNP) to convert 2-amino-6-mercapto-7-methylpurine (MESG) ribonucleoside to free ribose 1-phosphate and MESG, measuring MESG absorbance at 360 nm ($A_{360}$). Reactions in 100 µl volumes were performed in 96-well plates, with gentle shaking, and the $A_{360}$ measured every 5 min for 1 h, then every 30 min for an additional hour on a Biotek Synergy HT microplate reader at 28 °C.

For DGC measurement the pyrophosphate released from the diguanylate cyclase reaction is cleaved by addition of pyrophosphatase to liberate free Pi to react with PNP and MESG. For each reaction 10 µM of MBP-DcpA$_{Cyt}$ was added to a 100 µL reaction containing the following components: 24 mM Tris-HCl (pH 7.5), 5 mM $MgCl_2$, 45 mM NaCl, EnzChek reaction buffer, 1 µL PNP, 1 µL inorganic pyrophosphatase (previously diluted 10-fold in 1x reaction buffer), 20 µL MESG substrate, over a range of GTP concentrations, with volume adjusted to 100 µL with $ddH_2O$. Absorbance readings were normalized for spontaneous phosphate release by subtracting the $A_{360}$ reading of a control reaction containing c-di-GMP but no added protein.

*In vitro* PDE assays were also performed using the EnzChek Pyrophosphate Assay kit (Thermo Fisher, Waltham, MA). In this assay, linear pGpG produced as the product of the phosphodiesterase reaction served as substrate for calf intestinal alkaline phosphatase (CIP, New England Biolabs) to release 5′-Pi molecules that were detected using the MESG reaction. In the presence of free phosphate, 5 µM of MBP-DcpACyt was added to a 100 µL reaction containing the following components: 75 mM Tris-HCl (pH 7.5), 10 mM $MgCl_2$, 25 mM NaCl, 25 mM KCl, EnzChek reaction buffer, 1 µL PNP, 0.16 µL CIP (10,000 units/ml), 20 µL MESG substrate, and a range of c-di-GMP concentrations. Absorbance readings were normalized for spontaneous phosphate release by subtracting the $A_{360}$ reading of a control reaction containing GTP but no added protein.

**Rabbit-α-PruR and Rabbit-α-$DcpA_{peri}$ antibody production and purification.** $His_6$-PruR and Maltose-binding protein (MBP)-$DcpA_{Peri}$ fusions were expressed and purified as described in protein purification details and used by Josman, LLC for antibody production. A total of 5 mg of purified PruR (two doses at 2.5 mg each) or 0.75 mg of $DcpA_{peri}$ (one dose at 0.5 mg and a second at 0.25 mg) was injected into a rabbit host. Antibodies were purified from sera by incubating overnight at 4 $^{o}$C with either PruR or DcpA coupled Affigel-10 resin. This was then loaded onto a column and washed with Phosphate Buffered Saline (PBS) and eluted with 100 mM glycine (pH 2.5) and subsequently neutralized with 2 M Tris (pH 10.5). Elution fractions were pooled and dialyzed in PBS with 50% glycerol and then BSA was added to a final antibody concentration of 1 mg/mL. Antibodies were further purified to reduce non-specific binding. Briefly, nitrocellulose squares (~25 $mm^2$) were incubated with cell lysates generated from either a Δ*pruR* or Δ*dcpA* strain overnight at 4 $^{o}$C. Squares were then removed from the lysates and washed with PBS (three times for 5 min each) and blocked in 5% (w/v) Blotto solution (5% (w/v) non-fat dry milk in TBS-T) for 1 h at 4 $^{o}$C. Antibodies were then diluted 1:200 into PBS with 50% glycerol and incubated with the nitrocellulose squares overnight at 4 $^{o}$C. Finally, nitrocellulose squares were removed and antibodies were stored at -20 $^{o}$C.

**Western Blots.** Purified protein or whole-cell lysates ($OD_{600}$: 30 equivalent) were incubated at 90°C for 5 min with an equal volume of SDS sample loading buffer (60 mM Tris-HCl pH 6.8, 1% (w/v) 2-mercaptoethanol, 1% (w/v) SDS, 10% (v/v) glycerol, 0.01% (w/v)). 10 µL of denatured/lysed samples were separated by SDS-PAGE (10% pH 8.8 resolving, 5% pH 6.8 stacking) and electrophoresed (100 V ~1.5 h) using a discontinuous Tris-glycine Laemmli buffer system (11). Gels were incubated in transfer buffer (48 mM Tris-base (pH 8.3), 39 mM glycine, 1.3 mM SDS, 20% (v/v) methanol) for 20 min before transfer to nitrocellulose membrane using a Trans-Blot SD Semi-Dry Transfer Cell (Bio-Rad, Hercules, CA). Membranes were stained with Ponceau S staining solution (0.1% (w/v) Ponceau S, 5% (v/v) acetic acid) to ensure proper protein transfer. Membranes were washed for 5 min in Tris buffered saline (50 mM Tris (pH 7.6), 150 mM NaCl) before being blocked overnight rocking at 4°C in 5% (w/v) Blotto solution (5% (w/v) non-fat dry milk in TBS with 0.1% Tween-20 (TBS-T). Membranes were washed three times with TBS-T and incubated overnight rocking at 4°C with primary antibodies diluted 1:40,000 (Rabbit-α-PruR) or 1:20,000 (Rabbit-α-$DcpA_{peri}$) in 5% Blotto. Membranes were washed three times with TBS-T and incubated for 1 h at room temperature rocking with goat-α-rabbit secondary antibody conjugated to horseradish peroxidase (GAR-HRP) (Abcam, Cambridge, MA) diluted 1:20,000 in 5% Blotto. Membranes were washed with TBS-T and were developed with SuperSignal West Pico Luminol/Enhancer Solution (Thermo Fisher, Waltham, MA) and the HRP signal was detected using a Biorad ChemiDoc system using Image Lab software.

***A. tumefaciens* periplasmic fractionation.** *A. tumefaciens* periplasmic proteins were isolated using an adapted osmotic shock procedure developed by Dyé and Delmotte (12). Cultures were grown shaking overnight at 30 °C, and 5 mL of culture was centrifuged (11,750 *g* x 10 min., 4 °C). Cell pellets were resuspended in 1 mL cold osmotic shock buffer (100 mM Tris-HCl (pH 8.0), 1 M sucrose, 0.5 M ethylenediaminetetraacetic acid (EDTA), 1 mM phenylmethylsulphonyl fluoride (PMSF) and incubated on ice for 5 min. After incubation, cells were centrifuged (7,800 *g* x 3 min.) and cell pellets were warmed to room temperature. Cell pellets were then resuspended in 1 mL ice-cold $ddH_20$ and incubated on ice for 1 min, followed by addition of $MgCl_2$ to a final concentration of 1 mM. After 5 min incubation on ice, cells were centrifuged (13,200 *g* x 3 min.), and the supernatant (periplasmic fraction) was collected. Cell pellets (cytoplasmic fractions) were resuspended in 1 mL $ddH_20$ with 1 mM $MgCl_2$. Periplasmic and cytoplasmic fractions were separated by electrophoresis on SDS-PAGE and probed following the Western blot procedure described above.

***In vivo* DSS-Crosslinking.** Cells were grown to mid-exponential phase and after determining $OD_{600}$, two 2 ml samples were collected via centrifugation (9,000 *g* x 3 min) and each resuspended in 1 mL ice-cold PBS (20 mM sodium phosphate, 0.15 M NaCl; pH 8). 50 µL 15 mM disuccinimidyl suberate (DSS) in dimethyl sulfoxide (DMSO) was added to one sample (final concentration 0.75 mM DSS), and 50 µL DMSO was added to the second as a negative control. Samples were vortexed and rocked at room temperature for 30 min. The crosslinking reaction was quenched with the addition of 1 M Tris-HCl (pH 7.5) to a final concentration of 20 mM and then vortexed and rocked again at room temperature for 15 min. Cells were pelleted via centrifugation (9,000 *g* x 3 min). Cells were resuspended in SDS sample loading buffer to an $OD_{600}$ of 30, based on measurements prior to crosslinking.

***In vitro* analysis of pterin binding to PruR.** Purified $\Delta_{SS}$-PruR (50 µM in a 500 µl reaction volume) was combined with potential pterin and folate ligands (300 µM) in 100 mM potassium phosphate pH 6.0 + 5 mM 2-mercaptoethanol in an anaerobic chamber (Coy Laboratory Products, 97% $N_2$/3% $H_2$) at room temperature for 30 min. Folate, dihydrofolate ($H_2F$), tetrahydrofolate ($H_4F$), and D-neopterin were obtained from MilliporeSigma. Due to limited commercial availability 7,8-dihydromonapterin ($H_2MPt$) was prepared from L-xylose based upon the previously reported procedure (13). 7,8-Dihydroneopterin ($H_2NPt$) was obtained by alkaline zinc reduction of neopterin (14). The concentrations of all pterin stock solutions were determined by measuring their absorbance at pH 1.0 and using the reported extinction coefficients (14). To obtain $H_4MPt$ and $H_4NPt$, purified $His_6$-PruA (5 µM, expressed and purified as previously described; (15)) was combined with NADPH (1 mM) and the corresponding dihydropterins as substrates under anaerobic conditions at room temperature for 30 min, after which PruR was added to the PruA reaction mixture and incubated an additional 30 min. The samples were then applied to a small column (0.5 x 1 cm) of Ni-NTA resin (Cytiva) equilibrated with 50 mM HEPES, 300 mM NaCl, 20 mM imidazole (pH. 8.0) and washed with 2 mL of the same buffer. This step removed the $His_6$-PruA from the samples, which remained bound to the nickel resin. Next, the PruR-containing flow through and wash were combined (2.5 mL total) and applied to a PD-10 desalting column (Cytiva) that was previously equilibrated with 100 mM potassium phosphate (pH 6.0) + 5 mM 2-mercaptoethanol. The PruR protein was eluted with 3.5 mL of the same buffer. This step removed the unbound pterins present in the sample. The PruR sample was then concentrated to 0.5 mL under anaerobic conditions with an Amicon Ultra centrifugal filter unit (10 kDa molecular weight cutoff, MilliporeSigma). The protein was precipitated by addition of $CH_3CN$ (50% final v/v) and the sample was concentrated under vacuum to 100 µl. Since the reduced forms of pterins and folates are unstable and susceptible to air oxidation/degradation, the samples were chemically oxidized with iodine before HPLC analysis. Thus, 5 µl of an iodine solution (10% $I_2$, 20% KI in water) was added to the concentrated PruR sample. After 30 min, 5 µl of 1 M $Na_2S_2O_5$ was added to consume unreacted iodine. All PruR/pterin binding experiments were performed in triplicate and error bars are standard deviation.

**Analysis of pterins and folates by HPLC.** A Shimadzu HPLC system equipped with a RF-20 fluorescence detector and a Phenomenex Kinetex polar C18 column (150 x 4.6 mm) was utilized to analyze pterins and folates. Oxidized pterins are fluorescent (Excitation: 356 nm, Emission: 450 nm) while folate is not fluorescent and is instead observed by UV absorbance (283 nm). Solvent A was 0.1% HCOOH in water and Solvent B was 100% methanol. The flow rate was 0.7 mL/min and the column was equilibrated with 98% A/2% B. For pterins, the elution profile consisted of 2 min at 98% A/2% followed by an 18 min linear gradient to 60%A/40%B. For folates, the elution profile consisted of 2 min at 98% A/2% followed by an 18 min linear gradient to

30%A/70%B. All pterins and folates were analyzed in their fully oxidized forms (generated by iodine oxidation as described above).

**Molecular cloning and protein production for crystallization.** Putative *pruR* genes from *Agrobacterium tumefaciens* str. C58 (gi: AAK89898, residues 23-168), *Klebsiella pneumoniae* subsp. pneumoniae NTUH-K2044 (gi: BAH64245, residues 24-169), *Vibrio cholerae* O1 biovar El tor str. N16961 (gi: AAF95081, residues 22-156), and *Vibrio vulnificus* CMCP6 (gi: AAO08173, residues 19-153) were cloned using ligation independent cloning into vector pMCSG53 containing an N-terminal $His_6$-tag followed by a tobacco etch virus (TEV) protease cleavage site, encoding ampicillin resistance, and genes for rare codons (8, 16). The plasmid for *pruR* from *A. tumefaciens* was transformed into *E. coli* BL21-Gold (DE3) cells and the other plasmids into *E. coli* BL21 (DE3) (Magic) cells(2).

All proteins were expressed in M9 media (High Yield M9 SeMet media, Medicilon Inc.). The starting overnight culture was grown in Luria broth supplemented with 130 µg/ml ampicillin and 50 µg/ml kanamycin at temperature 37 °C and rotation 220 rpm. The next day, 3 L of M9 media supplemented with 200 µg/ml ampicillin and 50 µg/ml kanamycin were inoculated at 1:100 dilution with the overnight culture and incubated at temperature 37 °C and rotation at 220 rpm. Protein expression was induced at $OD_{600}$ of 1.8-2 by addition of 0.5 mM IPTG and the culture was further incubated at 25 °C, with shaking at 200 rpm for 18 hours (17) The cells were harvested by centrifugation at 8000 rpm for 20 min, resuspended (1 g of cells per 5 ml of lysis buffer(50 mM Tris (pH 8.3),

0.5 M NaCl, 10% glycerol, 0.1% IGEPAL CA-630) and frozen at −30 °C until purification.

Frozen suspensions were thawed and sonicated at 50% amplitude, in 5 s x 10 s cycle for 20 min on ice. The lysate was cleared by centrifugation at 18,000 *g* x 40 min at 4 °C, the supernatant was collected, and the protein was purified as previously described with some modifications (18). *N*-dodecyl-β-D-maltoside (DDM) was added to 1 mg/ml to the lysate from cells expressing PruR from *K. pneumoniae* prior to sonication. All proteins were purified using an ÅKTAxpress FPLC system (GE Healthcare). The supernatants were loaded into a His-Trap FF (Ni-NTA) column in loading buffer (10 mM Tris-HCl (pH 8.3), 0.5 M NaCl, 1 mM Tris (2-carboxyethyl) phosphine (TCEP), and 5% glycerol). The column was washed with 10 column volumes (cv) of loading buffer and 10 cv of washing buffer (10 mM Tris-HCl (pH 8.3), 1M NaCl, 25 mM imidazole, 5% glycerol). The proteins were eluted with elution buffer (10 mM Tris (pH 8.3), 0.5 M NaCl, 1 M imidazole), loaded onto a Superdex 200 26/600 column and separated in loading buffer. The protein was collected and analyzed by SDS-PAGE. The $His_6$-tag was cleaved by mixing 1:20 (protease:protein) with recombinant TEV protease overnight at room temperature during dialysis. PruR from *K. pneumoniae* precipitated after tag cleavage and was diluted in buffer, resuspended, and filtered. All cleaved proteins were separated from recombinant TEV protease, uncleaved protein and the $His_6$-tag peptide by Ni-NTA-affinity chromatography using loading buffer followed by loading buffer with 25 mM imidazole. The cleaved protein was collected in the flow-through in loading buffer, analyzed by SDS-PAGE for purity and $His_6$-tag cleavage. All proteins were concentrated to 7.2 – 12.5 mg/ml and set up for crystallization.

**Crystallization.** The proteins were set up for crystallization in buffer containing 10 mM Tris-HCl (pH 8.3), 150 mM NaCl and 1 mM TCEP as 2 µl crystallization drops (1 µl protein: 1 µl reservoir solution) in 96-well crystallization plates (Corning) using commercial Anions, Classics II, JCSG+, PACT, and PEG's II (QIAGEN) crystallization screens, in the absence or presence of 1 mM (for *A. tumefaciens*) or 2 mM (for all others) neopterin. Diffraction quality crystals of PruR from A. *tumefaciens* grew from the condition with 2.4 M sodium malonate (pH 7.0) and 2.1 M DL-malic acid (pH 7.0), 25% polyethylene glycol (PEG) 400, and were cryo-protected in 25% sucrose, 1.2 M sodium malonate (pH 7.0), and 1.5 M DL-malic acid (pH 7.0) respectively; the *K. pneumoniae* PruR crystals grew from 0.2 M lithium sulfate, 0.1 M Tris (pH 8.5), and 25% (w/v) PEG 3350 and the crystal was cryo-protected using the reservoir solution; the *V. cholerae* PruR crystals grew from 0.1 M Bis-Tris (pH 6.5), 28% (w/v) PEG monomethyl ether 2000 and the crystal was cryo-protected using the reservoir solution; the *V. vulnificus* PruR crystal grew from 0.1 M citric acid (pH 3.5), 3 M NaCl, and crystals were cryo-protected in 4.0 M sodium formate. Crystals were flash cooled in liquid nitrogen for data collection.

**Data collection, structure solution and refinement.** The data sets were collected at the beam lines 21ID-G and 21ID-F of the Life Sciences-Collaborative Access Team (LS-CAT) at the Advanced Photon Source (APS), Argonne National Laboratory. Images were indexed, integrated, and scaled using HKL-3000 (19). Structures of 7kom, 7rkb, and 7kp2 were solved in HKL-3000 using Automated Structure Solution package. Initial experimental phases were obtained from Selenium atoms of Se-Met using Single Anomalous Disperse (SAD) method. Structures of both crystals form for 7kos and 7kou were solved by Molecular Replacement using 7kom as the search model in PHASER. All models were refined using REFMAC (20). Manual corrections and visualization was done in Coot (21). Water molecules were generated automatically in ARP/wARP (22) and ligands were fit into maps using Coot. Translation-Libration-Screw (TLS) groups were generated using TLSMD server (23) and TLS corrections were applied at the final steps of the model refinement. Structures were validated using MolProbity (24) and coordinates of the model and experimental data were deposited to the Protein Data Bank with the assigned PDB codes 7kom, 7rkb, 7kp2, 7kos and 7kou. The search for structural homologs was done using DALI (25) structural alignments were done using FATCAT (26), and all Figs were prepared in PyMol (The PyMOL Molecular Graphics System, Version 2.0, Schrödinger, LLC).

#### Supplementary References.

1. S. L. Chiang, E. J. Rubin, Construction of a mariner-based transposon for epitope-tagging and genomic targeting. *Gene* **296**, 179-185 (2002).
2. K. Kwon, S. N. Peterson, "High-throughput cloning for biophysical applications in structural genomics and drug discovery: methods and protocols " in Structural Genomics and Drug Discovery, F. W. Anderson, Ed. (Springer New York, New York, NY, 2014), pp. 61-74.
3. F. Lassalle *et al.*, Genomic species are ecological species as revealed by comparative genomics in *Agrobacterium tumefaciens*. *Genome Biol Evol* **3**, 762-781 (2011).
4. N. Feirer *et al.*, A pterin-dependent signaling pathway regulates a dual-function diguanylate cyclase-phosphodiesterase controlling surface attachment in *Agrobacterium tumefaciens*. *mBio* **6**, e00156 (2015).

5. J. Xu *et al.*, Genetic analysis of *Agrobacterium tumefaciens* unipolar polysaccharide production reveals complex integrated control of the motile-to-sessile switch. *Mol Microbiol* **89**, 929-948 (2013).
6. S. R. Khan, J. Gaines, R. M. Roop, 2nd, S. K. Farrand, Broad-host-range expression vectors with tightly regulated promoters and their use to examine the influence of TraR and TraM expression on Ti plasmid quorum sensing. *Appl Environ Microbiol* **74**, 5053-5062 (2008).
7. E. R. Morton, C. Fuqua, Genetic manipulation of *Agrobacterium*. *Curr Protoc Microbiol* 10.1002/9780471729259.mc03d02s25, Unit 3D 2 (2012).
8. Y. Kim *et al.*, Chapter 3. High-throughput protein purification for x-ray crystallography and NMR. *Adv Protein Chem Struct Biol* **75**, 85-105 (2008).
9. J. Tempé, A. Petit, M. Holsters, M. Van Montagu, J. Schell, Thermosensitive step associated with transfer of the Ti plasmid during conjugation: possible relation to transformation in crown gall. *Proc. Natl. Acad. Sci. USA* **74**, 2848-2849 (1977).
10. S. R. Maloy, V. J. Stewart, R. K. Taylor, *Genetic analysis of pathogenic bacteria: a laboratory manual.* (Cold Spring Harbor Press, Plainview, NY, 1996).
11. S. R. Gallagher, One-dimensional SDS gel electrophoresis of proteins. *Current protocols in molecular biology / edited by Frederick M. Ausubel ... [et al.]* **Chapter 10**, Unit 10 12A (2006).
12. F. Dye, F. M. Delmotte, Purification of a protein from *Agrobacterium tumefaciens* strain A348 that binds phenolic compounds. *The Biochemical journal* **321 ( Pt 2)**, 319-324 (1997).
13. V. R. Sokya, W. Pfleiderer, R. Prewo, Pteridines: Synthesis and characteristics of 5,6-dihydro-6-( 1,2,3-trihydroxypropyl)pteridines: Covalent intramolecular adducts. *Helv. Chim. Acta.* **73**, 808-826 (1990).
14. R. L. Blakley, S. J. Benkovic, *Folates and Pterins: Chemistry and Biochemistry of Pterins* (John Wiley & Sons, Inc., 1985), vol. 2.
15. M. Labine *et al.*, Enzymatic and mutational analysis of the PruA pteridine reductase required for pterin-dependent control of biofilm formation in *Agrobacterium tumefaciens*. *J Bacteriol* 10.1128/JB.00098-20 (2020).
16. W. H. Eschenfeldt *et al.*, New LIC vectors for production of proteins from genes containing rare codons. *J Struct Funct Genomics* **14**, 135-144 (2013).
17. C. S. Millard *et al.*, A less laborious approach to the high-throughput production of recombinant proteins in *Escherichia coli* using 2-liter plastic bottles. *Protein Expr Purif* **29**, 311-320 (2003).
18. L. Shuvalova, Parallel protein purification. *Methods Mol Biol* **1140**, 137-143 (2014).
19. W. Minor, M. Cymborowski, Z. Otwinowski, M. Chruszcz, HKL-3000: the integration of data reduction and structure solution--from diffraction images to an initial model in minutes. *Acta Crystallogr D Biol Crystallogr* **62**, 859-866 (2006).
20. G. N. Murshudov *et al.*, REFMAC5 for the refinement of macromolecular crystal structures. *Acta Crystallogr D Biol Crystallogr* **67**, 355-367 (2011).
21. P. Emsley, K. Cowtan, Coot: model-building tools for molecular graphics. *Acta Crystallogr D Biol Crystallogr* **60**, 2126-2132 (2004).
22. R. J. Morris, A. Perrakis, V. S. Lamzin, ARP/wARP and automatic interpretation of protein electron density maps. *Methods Enzymol* **374**, 229-244 (2003).
23. J. Painter, E. A. Merritt, Optimal description of a protein structure in terms of multiple groups undergoing TLS motion. *Acta Crystallogr D Biol Crystallogr* **62**, 439-450 (2006).
24. V. B. Chen *et al.*, MolProbity: all-atom structure validation for macromolecular crystallography. *Acta Crystallogr D Biol Crystallogr* **66**, 12-21 (2010).
25. L. Holm, Dali server: structural unification of protein families. *Nucleic Acids Res* **50**, W210-W215 (2022).

26. Z. Li, L. Jaroszewski, M. Iyer, M. Sedova, A. Godzik, FATCAT 2.0: towards a better understanding of the structural diversity of proteins. *Nucleic Acids Res* **48**, W60-W64 (2020).

#### Supplemental Figure Legends

**Figure S1. PruR negatively regulates biofilm formation and is required to control both the DGC and PDE activity of DcpA. (A)** The pterin-dependent regulatory pathway model of biofilm formation in *A. tumefaciens*. PruA reduces $H_2$MPt to $H_4$MPt, the pterin then interacts with PruR, which regulates the activity of DcpA. Promotion of PDE activity leads to decreased cdGMP levels and decreased UPP biosynthesis, which controls biofilm formation. **(B, C):** Cultures of *A. tumefaciens* C58 WT or a ∆*pruR* mutant harboring either a vector control (pSRKGm), or this vector expressing $P_{lac}$-*pruR*, were tested for: **(B)** Congo Red staining by inoculating 3 μl spots on ATGN-CR plates supplemented with 75μg/mL Congo Red and 400 μM IPTG and incubated at 30°C for 48 h and **(C)** Biofilm formation. Assays performed in triplicate and error bars are standard deviation; *P* values calculated by comparing wild type to the empty vector strain and *pruR* complementation by standard two-tailed *t*-test. ***<0.001.

**Figure S2 *In vitro* pterin binding assays**. HPLC chromatograms of extracted and oxidized pterins bound to purified PruR. **(A)** MPt standard (10 μM) compared to PruR binding assays with **(B)** $H_2$MPt or **(C)** $H_4$MPT. **(D)** NPt standard (10 μM) compared to PruR binding assays with **(E)** $H_2$NPt or **(F)** $H_4$NPt. **(G)** Folate standard (2 μM) compared to PruR binding assays with **(H)** $H_2$F or **(I)** $H_4$F. **(J)** PruR control without added pterins. **(K)** BSA control with $H_2$MPt and PruA/NADPH. PruR (50 μM) was incubated with $H_2$MPt or $H_2$NPt in the presence or absence of pteridine reductase PruA to assess binding of tetrahydro vs dihydro pterins, respectively. Binding experiments were also performed with commercially available $H_2F$ and $H_4F$. Chromatograms are representative reactions for triplicate assays. The chromatograms are injections from concentrated 100 µl samples while the original binding experiments were performed in a 500 µl reaction volume. The concentrations reported in Figure 1 of the main manuscript are the extrapolated amounts of pterins bound to PruR in the original 500 µl reaction volume.

**Figure S3. DcpA has diguanylate cyclase and phosphodiesterase activity *in vitro*.** Enzymatic activity of MBP-DcpA$_{cyt}$ **(A)** DGC and **(B)** PDE activity. Increasing concentrations of the substrates GTP or cdGMP were added as indicated, respectively, and DGC and PDE activity were determined indirectly by release of inorganic phosphate (Pi) in a coupled enzyme assay in which purine nucleoside phosphorylase (PNP) utilizes the Pi to convert 2-amino-6-mercapto-7-methylpurine (MESG) riboside to free MESG, measuring its absorbance at 360 nm ($A_{360}$).

**Figure S4. Examination of PruR signal sequence and potential for lipidation. (A)** Alkaline phosphatase assay in *A. tumefaciens* C58 WT or the Δ*pruR* mutant containing $P_{lac}$ expression plasmids harboring *pruR* without *phoA*, full-length *pruR* fused at its C-terminus to the 5'-end of *phoA*, or the codons for the secretion signal of *pruR* fused to *phoA* (*pruR*$_{SS}$-*phoA*). Activity in Miller Units. Assays performed in triplicate and error bars are standard deviation. **(B)** Ectopic expression of PruR-Cys$_{19}$ mutated to an alanine (*pruR*$_{C19A}$) was tested for complementation in biofilm assays. The ratio of acetic acid-solubilized CV absorbance ($A_{600}$) from 48 h biofilm assays normalized to the $OD_{600}$ planktonic turbidity of the same culture. Assays performed in triplicate and error bars are standard deviation; *P* values calculated by standard two-tailed *t*-test compared to wild type. (*P* values, *** <0.001,). **(C):** Plasmid-borne mutant *pruR* alleles (expressed with *dcpA*) with the N-terminal secretion signal replaced with that for MalE or DsbA from *E. coli* were ectopically expressed from $P_{lac}$ in the Δ*pruR* mutant. Cultures were induced with 50 μM and 75 μM IPTG, and periplasmic fractionation was performed. Western blot of cytoplasm/membrane (C/M) and periplasmic fractions using α-PruR antibody. Antibody binding detected with GAR-HRP secondary antibody and a chemiluminescent substrate on a BioRad ChemiDoc. Pro - Pro-peptide; Mat - Mature protein **(D)** Congo Red assays of the indicated strains. 3 μl spots were inoculated on ATGN-CR plates supplemented with 75 μg/mL Congo Red and 400 μM IPTG and incubated at 30°C for 48 h.

**Figure S5. Reciprocal detection of the PruR-DcpA complex using antibody against DcpA$_{peri}$.** Western blot probing for DcpA following i*n vivo* crosslinking with DSS (0.75 mM) in *A. tumefaciens* C58 wild type, a strain ectopically expressing $P_{lac}$-*pruR-dcpA*, a strain ectopically expressing $P_{lac}$-*pruA* (both with 400 μM IPTG) or a strain lacking *dcpA* (same extracts as in Fig. 3). Black arrows, PruR-DcpA, 87 kDa, DcpA, 71 kDa. α-DcpA$_{Peri}$ used at 1:20,000 dilution and antibody binding detected with GAR-HRP secondary antibody and chemiluminescent substrate on a BioRad ChemiDoc. Non-specific bands serve as protein loading controls.

**Figure S6. Multiple amino acid sequence alignments for PruR and DcpA homologs. (A)** Sequence alignments of PruR and its homologs. N-terminal signal sequence (Sec-Sig) is bracketed. Asterisks label residues conserved among SUOX family proteins, and the arrow indicates the position in canonical SUOX proteins at which a cysteine residue conjugates the molybdenum of the MoCo cofactor. Aligned PruR homologs, *Agrobacterium vitis* S4; AVI_RS19925, *Ochrobactrum anthropi* ATCC 49188, OANT_RS16195; *Klebsiella oxytoca* NCTC13727, EL226_RS11270; *Klebsiella pneumoniae* pneumoniae KPNIH16, KPNIH16_RS0129315; *Raoultella terrigena* NCTC13098, ELY08_RS09705; *Enterobacter* sp. HMSC055A11 HMPREF2540_RS13530; *Enterobacter cloacae* cloacae ENHKU01; ECENHK_RS04895; *Shigella flexneri* 1235-66; SF123566_7916; *Vibrio cholerae O1* biovar El Tor str. N16961, VC1933 *Vibrio vulnificus* CMCP6, VV1_RS2110; *Pseudomonas aeruginosa* PA01, PA2869. **(B)** Periplasmic domains of DcpA and homologs.  Similar highlighting scheme as in Panel A. Arrows indicate the two fully conserved residues. Aligned DcpA homologs; *O. anthropi* ATCC 49188, OANT_RS16200; *Klebsiella pneumoniae* pneumoniae KPNIH16, KPNIH16_RS02140; *Raoultella terrigena* NCTC13098, ELY08_RS09710; *Pseudomonas aeruginosa* PA01, PA2870, *Vibrio cholerae O1* biovar El Tor str. N16961, VC1934, *Vibrio vulnificus* CMCP6, VV1_RS21105. Yellow highlighting, fully conserved; Light blue highlighting, partial conservation; Green highlighting, similar R-group chemistry.

**Figure S7. PruR pterin binding pocket occupied by adjacent protomer in the crystal structure.** Space-filling model of *A. tumefaciens* (7kos). Q152 of neighboring molecule in the crystal structure, which is projected into the pterin binding pocket, is shown as a stick model with carbons in yellow, oxygens in red and nitrogens in blue.

Q152 is a part of the short α-helix (α7), which is located between β9 and β10, and it is shown in yellow.

**Figure S8. Pterin binding pocket of PruR. (A,B )** Two alternative binding modes of dihydroneopterin in the structure of PruR from *V. cholerae* (7kp2). **(C)** The structure of PruR from *K. pneumoniae* (7rkp) with neopterin bound to the pocket. The binding pocket is represented as an electrostatic surface potential and neopterin as a stick model. Carbons are in yellow (alternative conformation A, 7kp2), grey (alternative conformation B, 7kp2) and green (7rkb), oxygens in red and nitrogens in blue.

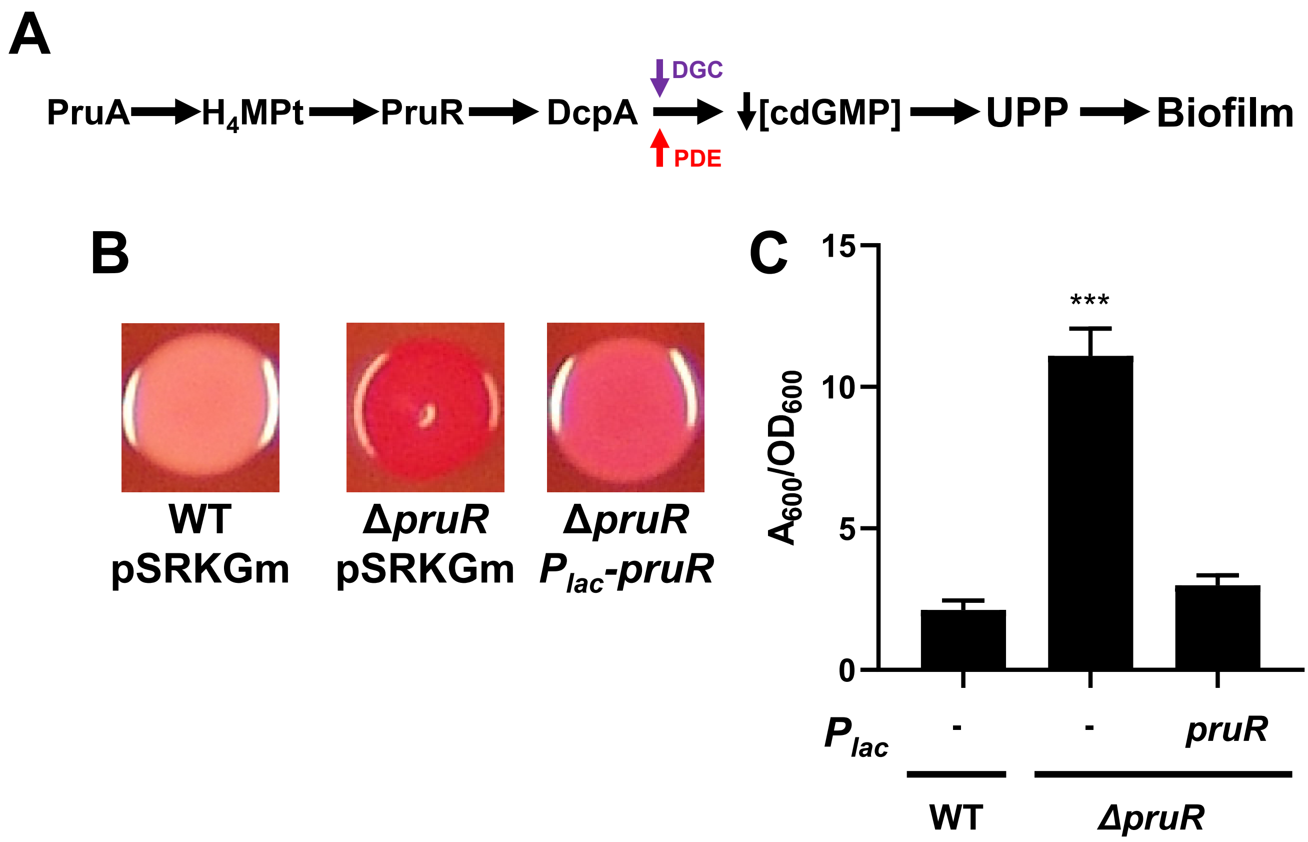

**Figure S1 – PruR-DcpA**

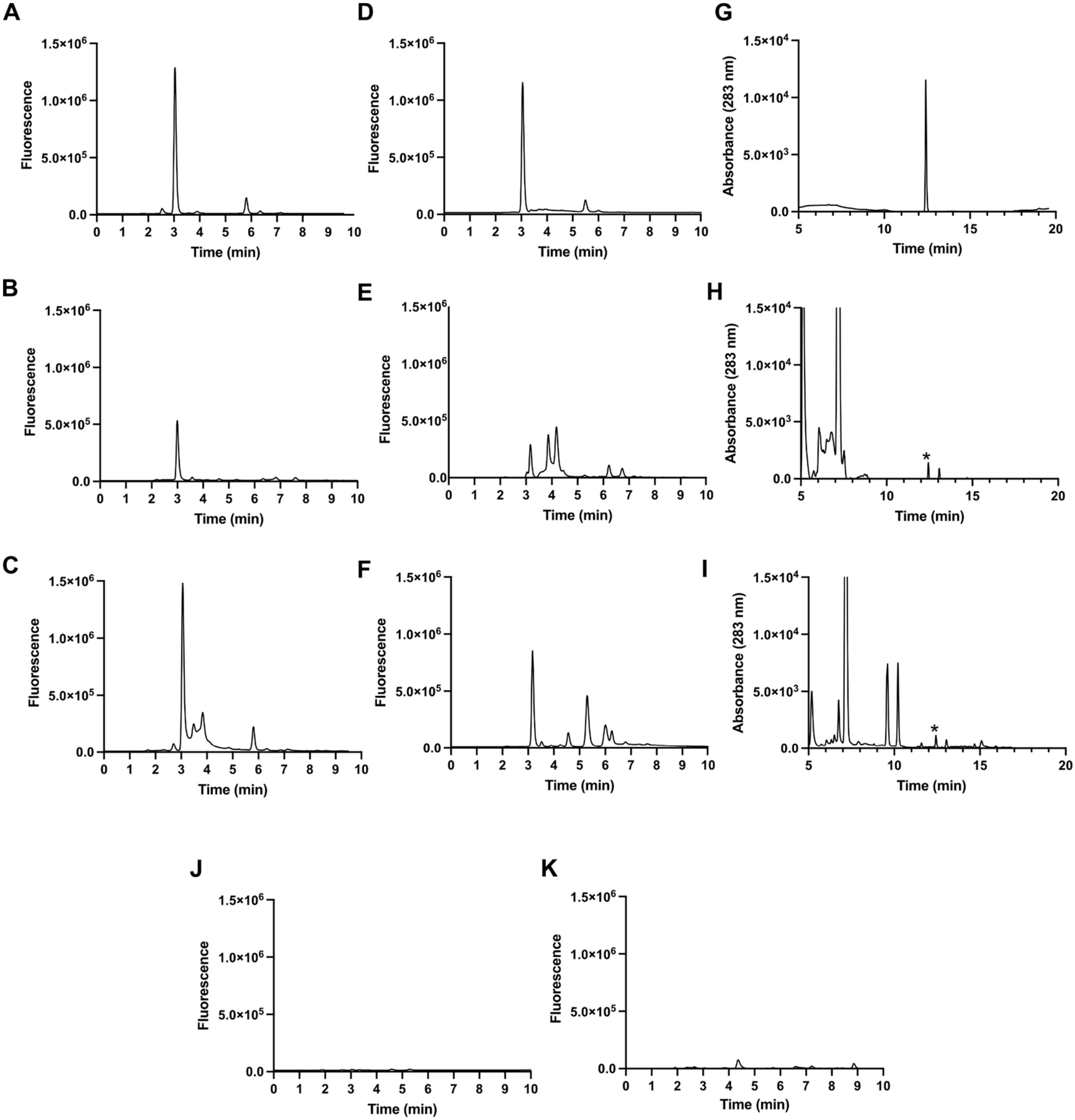

### Figure S2 - PruR-DcpA

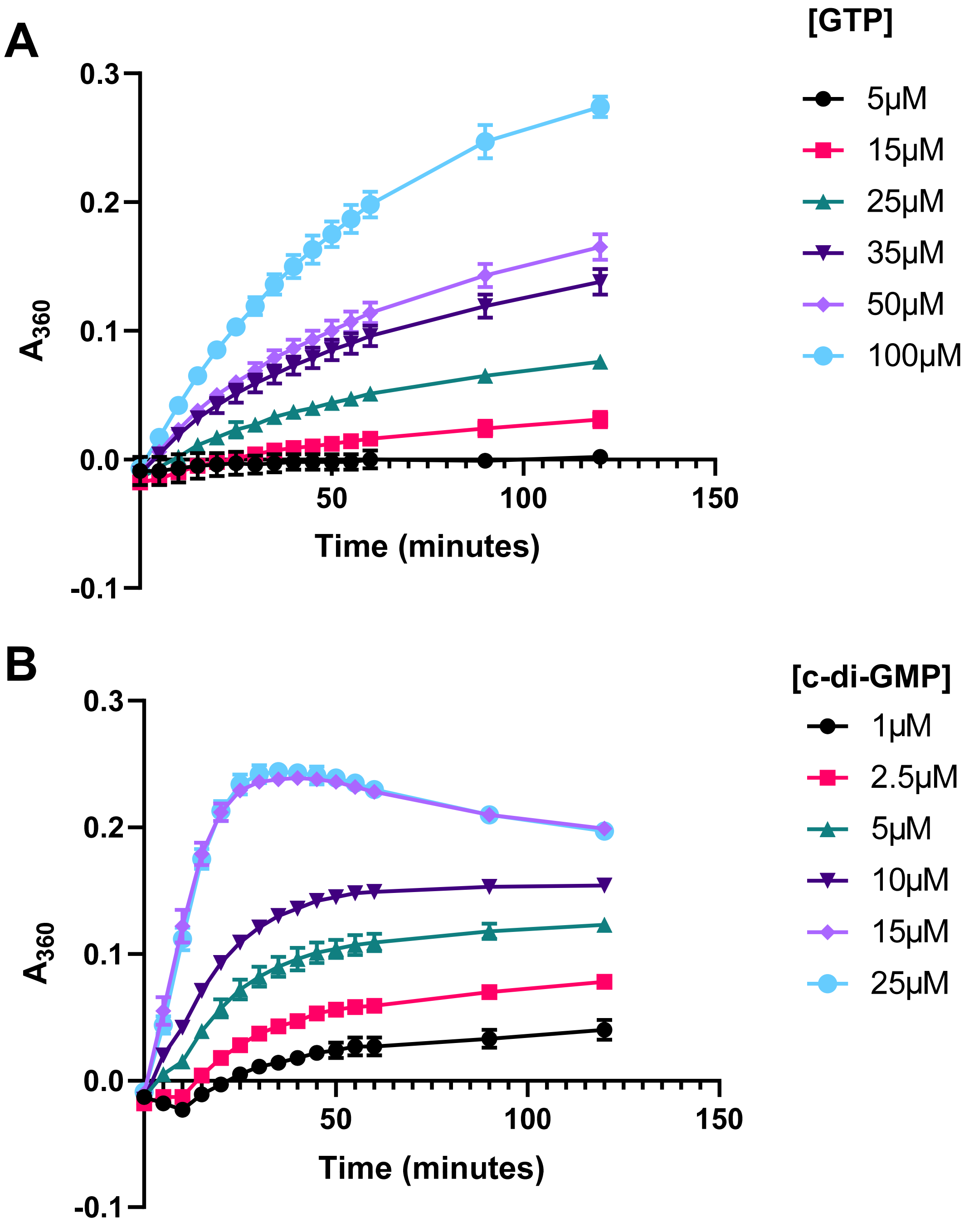

**Figure S3 - PruR-DcpA**

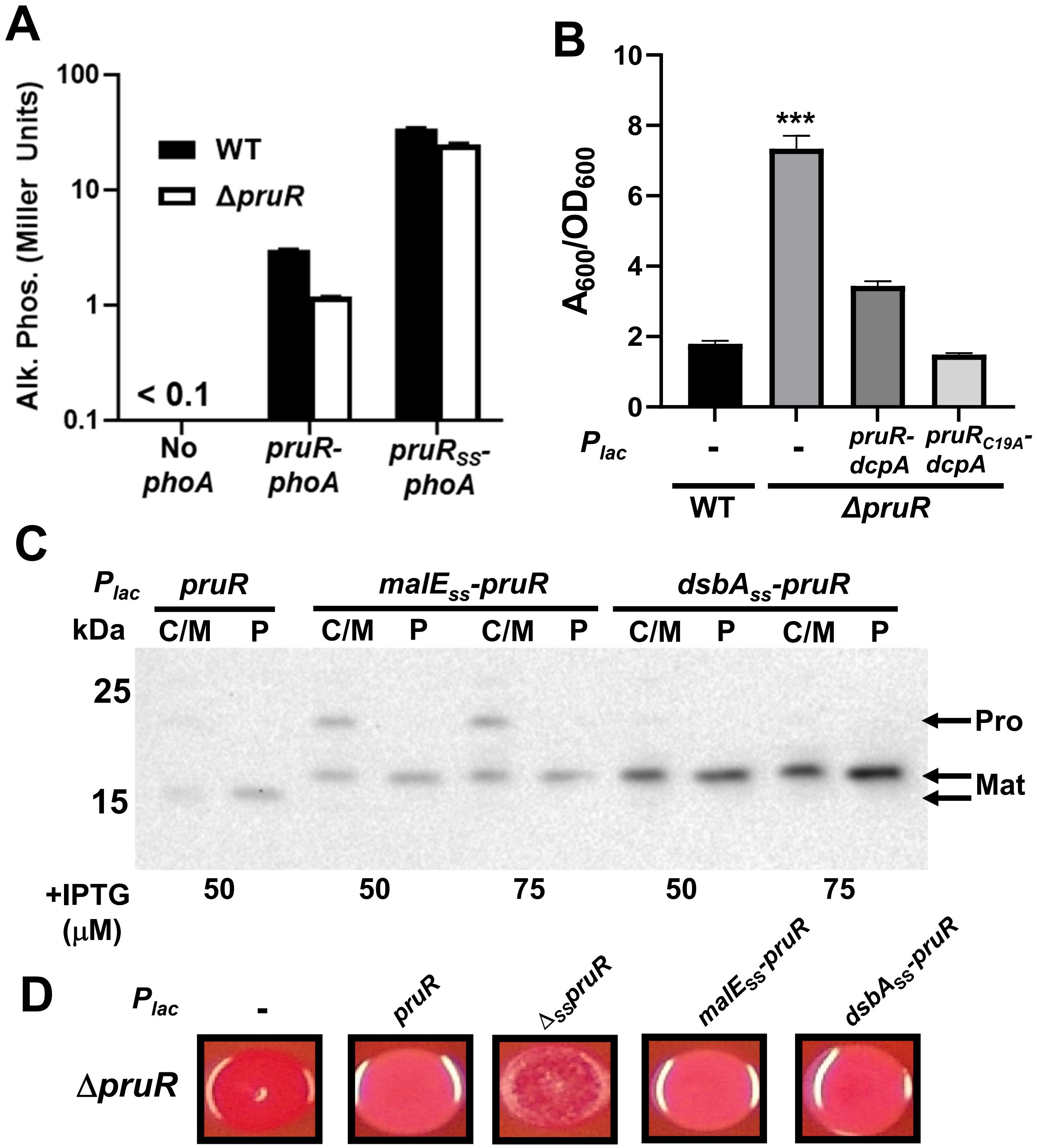

**Figure S4 – PruR-DcpA**

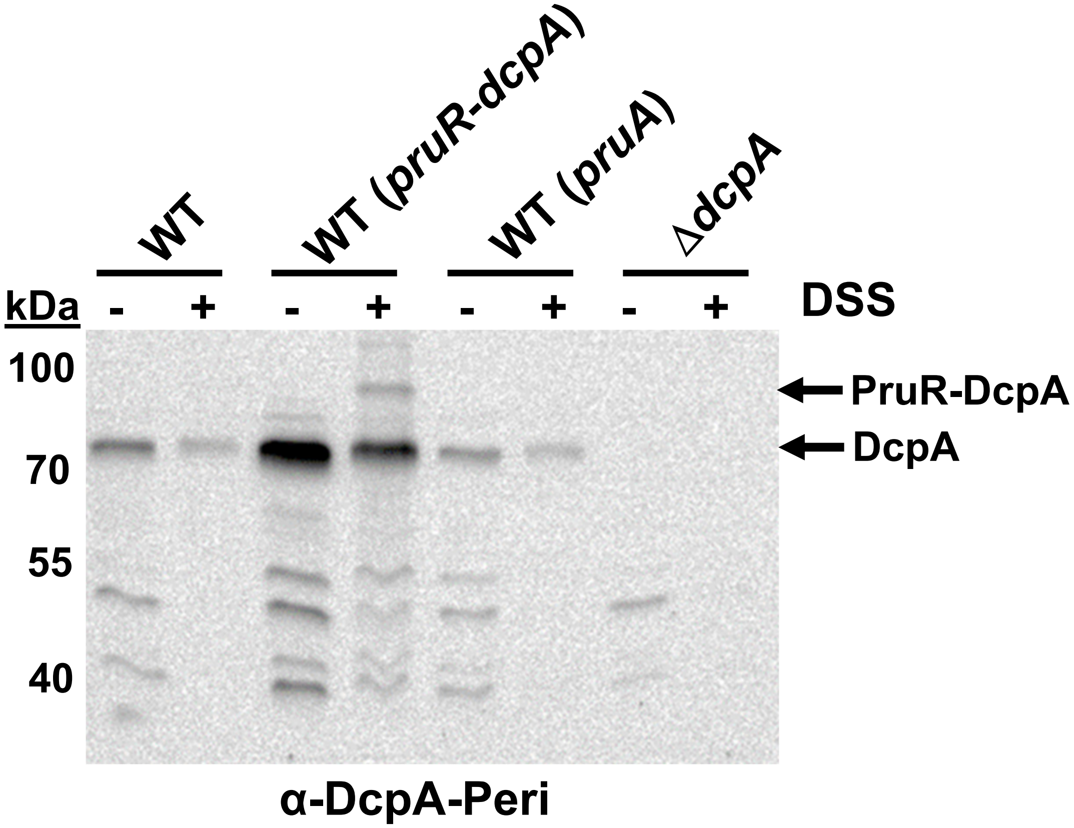

**Figure S5 – PruR-DcpA**

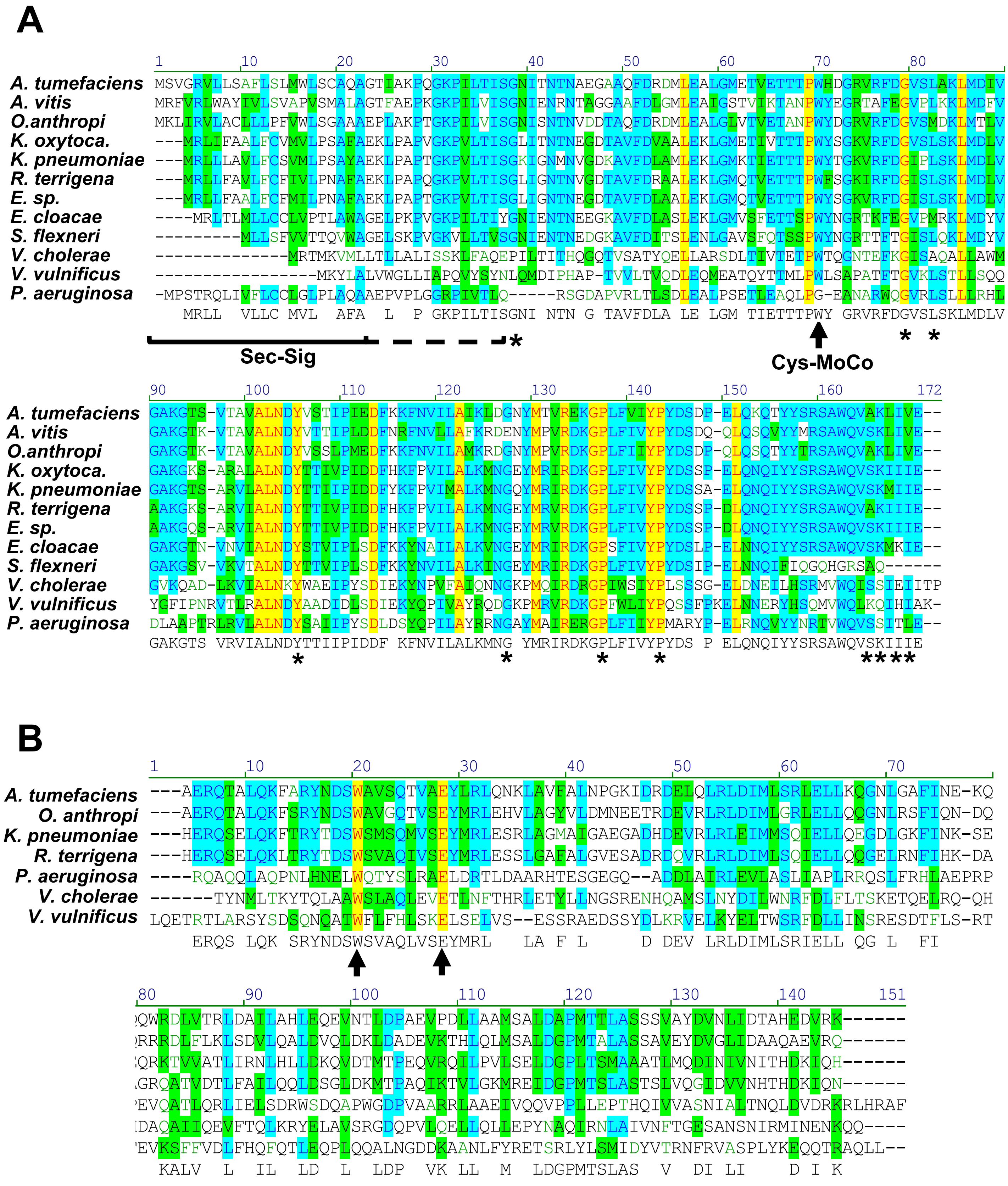

**Figure S6 – PruR-DcpA**

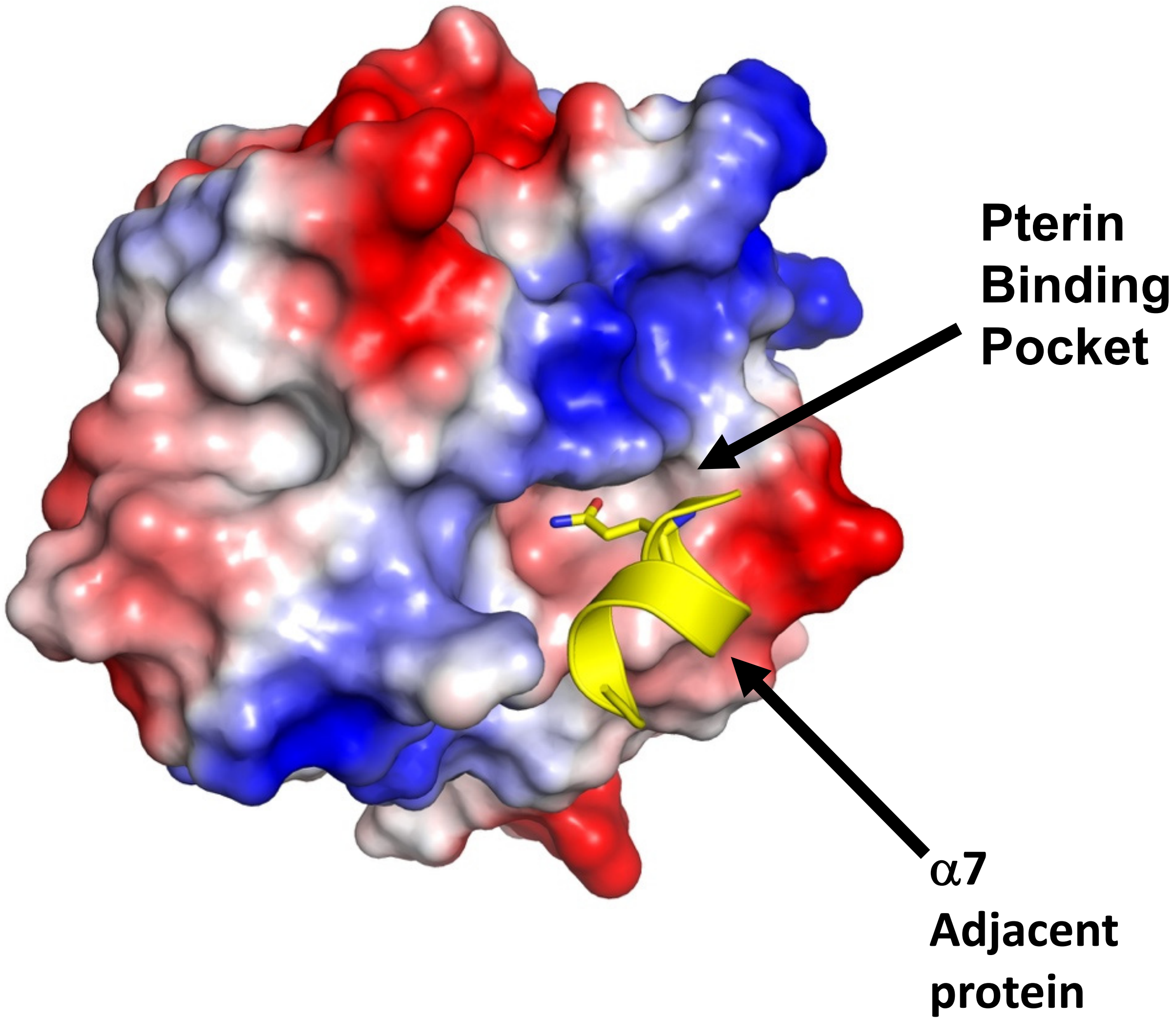

**Figure S7 - PruR-DcpA**

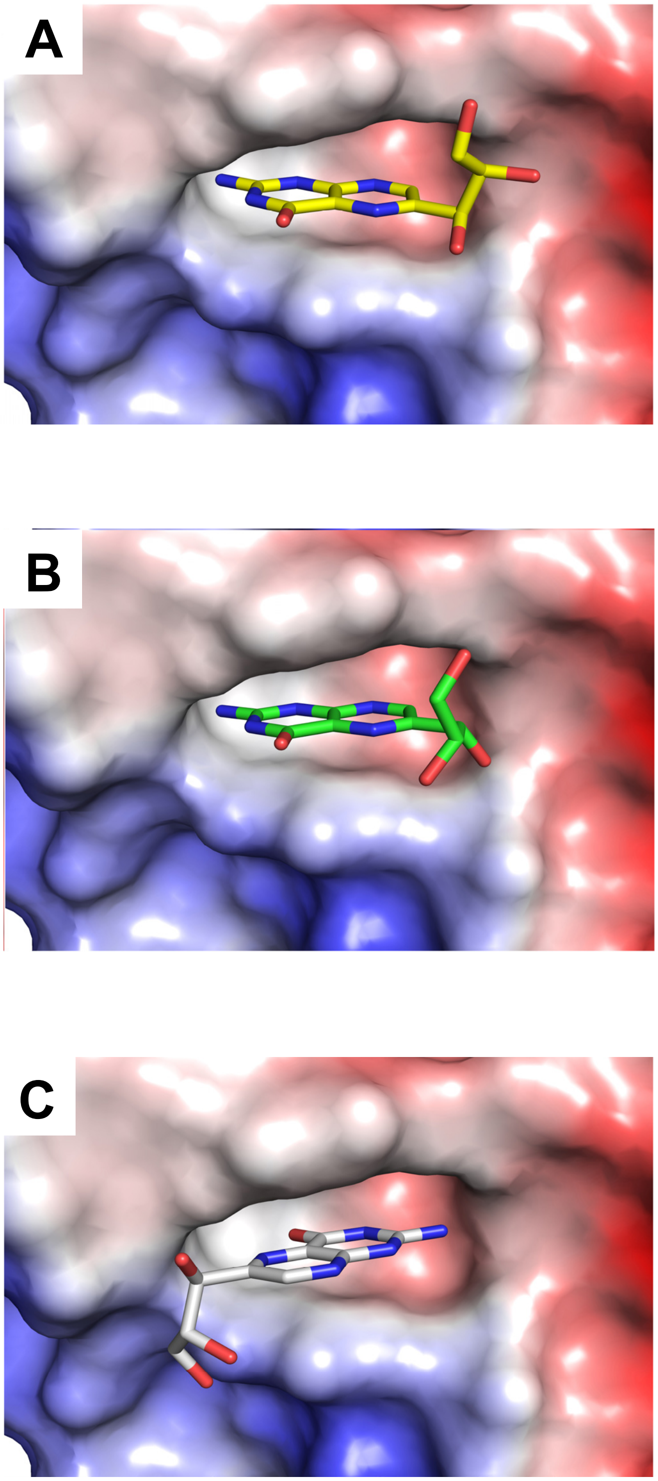

**Figure S8. Pterin binding pocket of PruR.**
